# Mitochondrial dsRNA: A Hidden Source of Immunogenic RNA in ADAR1 Deficiency

**DOI:** 10.64898/2026.01.06.698045

**Authors:** Haoqing Shen, Vincent Tano, Jian Han, Vanessa Hui En Ng, Wei Liang Gan, Sze Jing Tang, Ethan Tan, Shu Qin Peng, Larry Ng, Bryan Y. L. Ng, Ka Wai Leong, Xiaohui Sui, Jesslyn Zhou, Leilei Chen

**Affiliations:** Cancer Science Institute of Singapore, National University of Singapore; Singapore 117599, Singapore; NUS Center for Cancer Research, Yong Loo Lin School of Medicine, National University Singapore; Singapore 117594, Singapore; Department of Anatomy, Yong Loo Lin School of Medicine, National University of Singapore; Singapore 117594, Singapore

## Abstract

Double-stranded RNA (dsRNA) triggers immune responses during viral infections, but self-derived dsRNA can activate similar pathways. To prevent this, the body relies on mechanisms like ADAR1, an RNA-editing enzyme essential for immune regulation. Dysfunction of ADAR1 is linked to various diseases, yet the nature and role of dsRNAs accumulating in its absence remain unclear. Here, we identify mitochondrial dsRNA (mt-dsRNA), transcribed from the mitochondrial genome, as a major contributor to the endogenous dsRNA pool in ADAR1-deficient human and murine cells. We propose a "Draw-and-Release" model, where ADAR1 loss increases mitochondrial reactive oxygen species (mtROS), causing mt-dsRNA accumulation in the mitochondrial matrix ("Draw" phase) and its immune-activating release into the cytosol upon mitochondrial protein dysfunction ("Release" phase). This study highlights the importance of mitochondrial integrity in mitigating ADAR1-related pathologies.

## Main Text

The cellular innate immune response to viral infection is typically activated when viral double-stranded RNA (dsRNA) is recognized by host dsRNA sensors. Recent studies suggest that mammalian cells can also accumulate self-derived dsRNA, which may activate type-I interferon (IFN) response in a manner similar to viral infections. These self-derived dsRNAs can originate from various sources, such as genomic repeats (*1*), mitochondrial dsRNAs (mt-dsRNAs) (*2*), retained introns (*3*), and RNA modifications (*4–6*). Given the diverse sources of self-derived dsRNA, one key challenge for immune surveillance lies in preventing excessive accumulation of dsRNA in the cytoplasm, where the dsRNA sensors localize. In vertebrates, various mechanisms have evolved to ensure immune tolerance towards self-derived dsRNAs, distinguishing between foreign and self RNA ligands. Adenosine-to-inosine (A-to-I) RNA editing, catalyzed by ADAR (adenosine deaminase that acts on RNAs) proteins, has recently been identified as one of the most critical mechanisms. This editing process can mark cellular dsRNA as “self”, altering the Watson-Crick A-U base pairs to weaker I.U wobble pairs that destabilize dsRNA and prevent sensing by the melanoma differentiation-associated protein 5 (MDA5)–mitochondrial antiviral signalling protein (MAVS) pathway (*7–11*). Individuals with loss-of-function mutations of ADAR1 often develop Aicardi-Goutières syndrome (AGS), an autoimmune disease characterized by spontaneous IFN production (*12*). Multiple lines of evidence have shown that ADAR deficiency triggers accumulation of cellular dsRNAs, particularly in the cytoplasm, across different cell types in both humans and mice (*13–17*). However, the specific types of accumulated dsRNA, their immunogenic properties, and their primary contributions to the immune response in ADAR-deficient cells has yet to be thoroughly investigated. Inverted repeat *Alu* (IR*Alu*) elements embedded in the 3’ untranslated regions (3’UTRs) of mRNAs have long been considered the primary and abundant source of immunogenic dsRNA arising from ADAR1 loss. However, studies in both mice and human cells suggested that only a subset of unedited cellular dsRNAs are truly immunogenic (*18, 19*). Notably, emerging evidence also highlights rare *cis*-natural antisense transcripts (*cis*NATs) as critical contributors to immunogenicity (*20*). Moreover, a global RNA secondary structure analysis also revealed that A-to-I editing stabilizes more RNA duplexes than it destabilizes (*21*), challenging the prevailing notion that A-to-I (A-U to I·U) editing primarily inactivates RNA sensing. These findings collectively raise the question of the hidden sources of cellular dsRNA in the cytoplasm following ADAR1 deficiency and the precisely controlled mechanisms that restrict their immunogenicity to maintain immune homeostasis.

To address these questions, we conducted a genome-wide CRISPR screen to identify potential immune protectors involved in dsRNA sensing and innate immune response. Unexpectedly, mitochondrial proteins emerged as top candidates, and mt-dsRNAs, transcribed from the mitochondrial genome, accounted for a substantial portion of the endogenous dsRNA pool in both human and murine cells following ADAR1 deficiency. Through comprehensive functional and mechanistic investigations, we suggested a “*Draw*-and-*Release*” model in which the loss of ADAR1, independent of its catalytic function, leads to elevated mitochondrial reactive oxygen species (mtROS), causing the accumulation of mt-dsRNAs within the mitochondrial matrix. However, these mt-dsRNAs remain sequestered in the mitochondria during the “*Draw*” phase. Their release into the cytosol, triggering an immune response, only occurs during the “*Release*” phase, when the mitochondrial proteins (e.g., SOD2) identified in our CRISPR screen are defective or downregulated. We validated these findings both *in vitro* across multiple human and mouse cell lines and *in vivo* using liver-specific *Adar* conditional knockout (KO) mouse models. We demonstrated that reducing mtROS attenuates the cytosolic release of mt-dsRNAs in hepatocytes, reducing hepatic inflammation and tissue damage caused by ADAR1 loss. While ADAR1-mediated RNA editing has traditionally been linked to destabilizing IR*Alu*s-derived dsRNAs to prevent immune activation, our study uncovers an editing-independent role of ADAR1 in suppressing mt-dsRNA production and accumulation. This, in turn, reduces the risk of their release into the cytosol and the subsequent triggering of innate immune activation. These findings reveal a previously unrecognized role of mt-dsRNAs as immune-activating RNA ligands in the context of ADAR1 dysfunction and highlight the importance of maintaining mitochondrial integrity and function to mitigate pathologies associated with ADAR1 deficiency.

## Results

### Mitochondrial proteins are identified as top immune protectors in dsRNA sensing and innate immune response

Recent studies including ours reported several positive regulators of IFN response to self dsRNAs upon ADAR1 loss, such as *GRN*, *GGNBP2*, *CNOT10*, and *CNOT11 ^ref^* (*16, 22*). In this study, we intended to gain a comprehensive understanding of the dynamic regulation of self-dsRNA sensing by conducting a genome-wide CRISPR screen to identify genes that negatively control dsRNA sensing and IFN response following ADAR1 loss. We first generated ADAR1-knockout (ADAR1^KO^) HEK293T cells, which are known and validated to exhibit insensitivity to ADAR1 loss due to its high tolerance to cytoplasmic dsRNA accumulation (**fig. S1, A-C**) (*5, 23*). However, this cell model retains an intact dsRNA sensing machinery, as evidenced by detectable levels of key components, including MDA5, RIG-I, PKR, MAVS, IRF3, and TBK1, among others (**fig. S1D**), as well as significant *IFNB1* induction upon treatment with the dsRNA analogue polyinosinic-polycytidylic acid (polyIC) (**fig. S1E**).

We constructed a lentiviral reporter construct contains an EGFP gene driven by a 1,500 bp promoter region of the human *IFNB1* gene, enabling real-time quantification of initial IFN production and immune activation in response to dsRNA stimuli (**Fig. 1A** and **fig. S1, f and g**). ADAR1^KO^ and non-targeting (NT) control HEK293T cells were transduced with the reporter construct and then subjected to the CRISPR screen. ‘IFN-high’ and ‘IFN-low’ cells were sorted and sent for sgRNA sequencing (**Fig. 1A**). Using the MAGeCK bioinformatic pipeline (*24*), we calculated the enrichment of each sgRNA in the ‘IFN-high’ *versus* ‘IFN-low’ populations for each cell line (FDR < 0.05 and |log_2_(fold change, FC)| > 1). A total of 28 and 6 genes were identified as enriched in IFN-high ADAR1^KO^ and NT cells, respectively, with 4 genes overlapping between the two cell groups (**Fig. 1B, fig. S1H,** and **data S1**). Among these hits, there are previously identified immune suppressors, such as TRAIP (*25*), LSM11 ^ref^ (*26*), and COPA (*27*) (listed in the *top-right* corners of **Fig. 1B** and **fig. S1H**). Unexpectedly, nearly half of the hits identified in ADAR1^KO^ cells are nuclear genes encoding mitochondrial proteins (hereafter referred to as “mito-genes”) (listed in *top-left* corner, **Fig. 1B** and **Fig. 1C**), suggesting a critical role for mitochondria in controlling dsRNA sensing and innate immune response upon ADAR1 loss. Notably, 7 out of the top 10 hits identified in ADAR1^KO^ cells belong to mito-genes, while 6 of them are exclusive in ADAR1^KO^ cells. Silencing each of these 6 mito-genes in ADAR1^KO^ cells expressing the GFP-based reporter led to a significant upregulation of GFP expression and *IFNB1* transcripts, which was not observed in control siRNA (siNC)-treated ADAR1^KO^ cells or in any group of NT cells (**Fig. 1D**). Altogether, these findings highlight a previously unrecognized link between mitochondrial function and innate immunity in the context of ADAR1 deficiency.

**Fig. 1.**
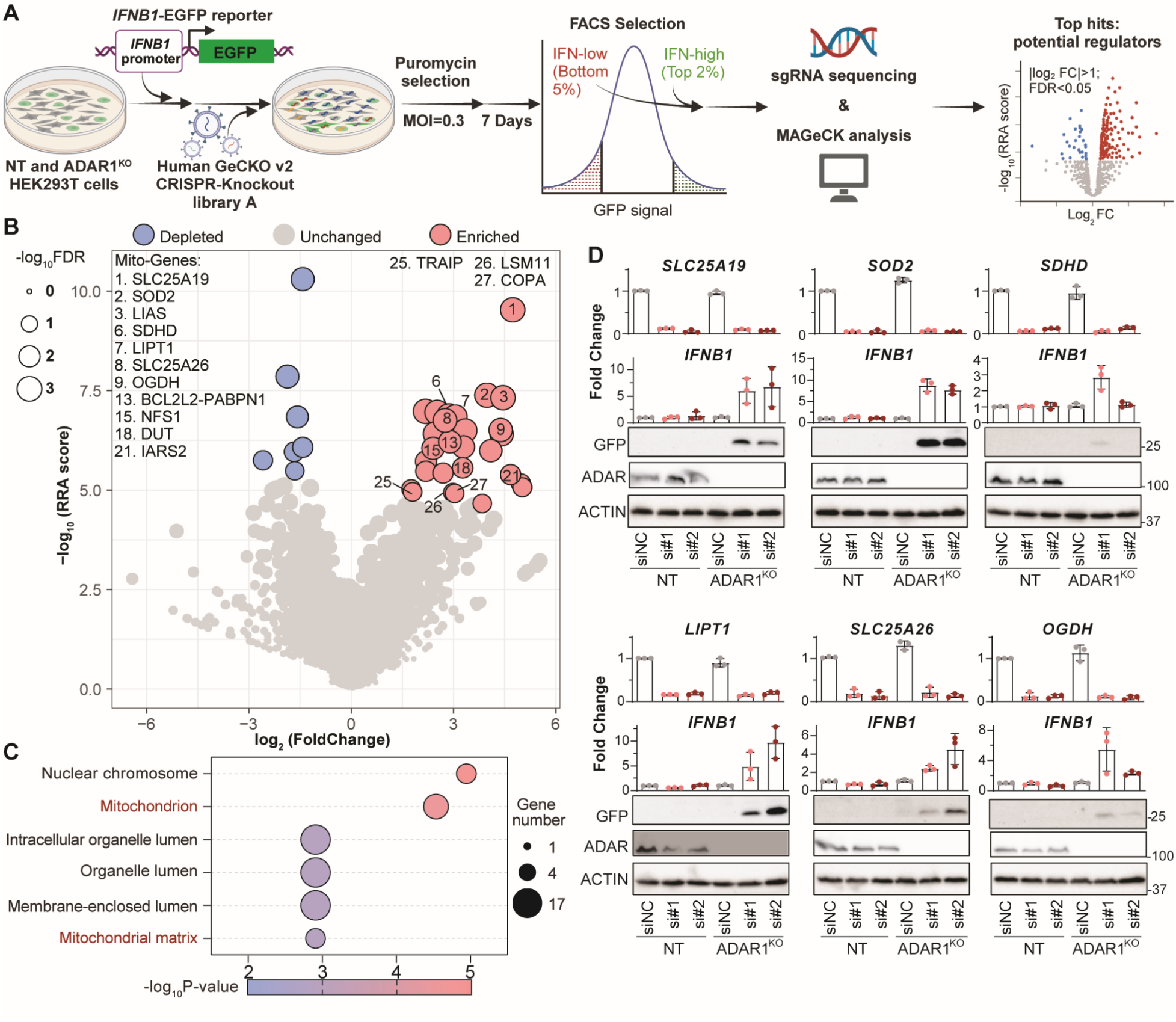
A genome-wide screen identified a subset of mitochondria proteins as potential negative regulators of IFN response. (**A**) Workflow of the phenotypic CRISPR screening. GFP fluorescence was measured by flow cytometry to quantify the level of immune response activation and cells with the highest (top 2%; ‘IFN-high’) and lowest (bottom 5%; ‘IFN-low’) GFP signals were sorted and collected for sgRNA sequencing. MAGeCK bioinformatic pipeline (*24*) was used to identify potential negative regulators of IFN response. Two independent screens were performed. Created in BioRender. Shen, H. (2025) https://BioRender.com/a69p174. (**B**) Volcano plots illustrating the top hits identified from the CRISPR screen conducted in ADAR1^KO^ HEK293T cells. Each dot represents a gene, with *x*-axis displaying the log_2_(Fold change) calculated based on the enrichment of sgRNAs targeting the indicated gene in ‘IFN-high’ population versus ‘IFN-low’ population, and *y*-axis showing the -log_10_ Robust Rank Aggregation (RRA) score. Significance was determined using thresholds of FDR < 0.05 and |log_2_(Fold change)| > 1, as indicated by dot size. Genes significantly enriched in the ‘IFN-high’ population are highlighted as red dots. Mitochondrial protein-coding genes (mito-genes) and previously reported innate immune repressors are listed in the upper-left and upper-right corners, respectively. The numbers preceding the gene names represent their respective rankings. (**C**) Gene ontology (GO) analysis of genes enriched in the ‘IFN-high’ population in ADAR1^KO^ cells. Dot size corresponds to the number of genes with each GO category, while the *x*-axis and dot color indicate the -log_10_(p-value), reflecting the statistical significance of the enrichment. (**D**) Quantitative PCR (qPCR) analysis of expression changes in identified mito-genes and *IFNB1*, along with western blotting (WB) of GFP and ADAR1 expression levels, following treatment with siRNAs targeting the indicated mito-genes in both NT and ADAR1 ^KO^ HEK293T cells. Two siRNAs were used for each target gene. Each dot represents the mean value of technical triplicates from an independent experiment. Data are presented as the mean ± S.D. of 3 independent experiments (biological replicates). siNC, non-targeting siRNA.

### ADAR1 deficiency leads to the accumulation of dsRNAs in the mitochondrial matrix

The role of mitochondrial proteins in regulating the innate immune response triggered by ADAR1 loss raises questions about the subcellular localization of potential immunogenic dsRNAs. Notably, in ADAR1^KO^ HEK293T cells, elevated dsRNAs formed granules that predominantly colocalized with mitochondria (**Fig. 2, A and C**), and RNase III treatment confirmed that these granules indeed represent dsRNA (**fig. S2A**). Similar results were obtained in other cell lines, including the murine hepatocyte cell line AML12 (**Fig. 2, B and C,** and **fig. S2B)**, as well as the human esophageal squamous cell carcinoma cell line EC109 and the murine hepatoma cell line Hepa1-6 (**fig. S2, C and D**), suggesting that our findings are not restricted to a specific human or murine cell line.

**Fig. 2.**
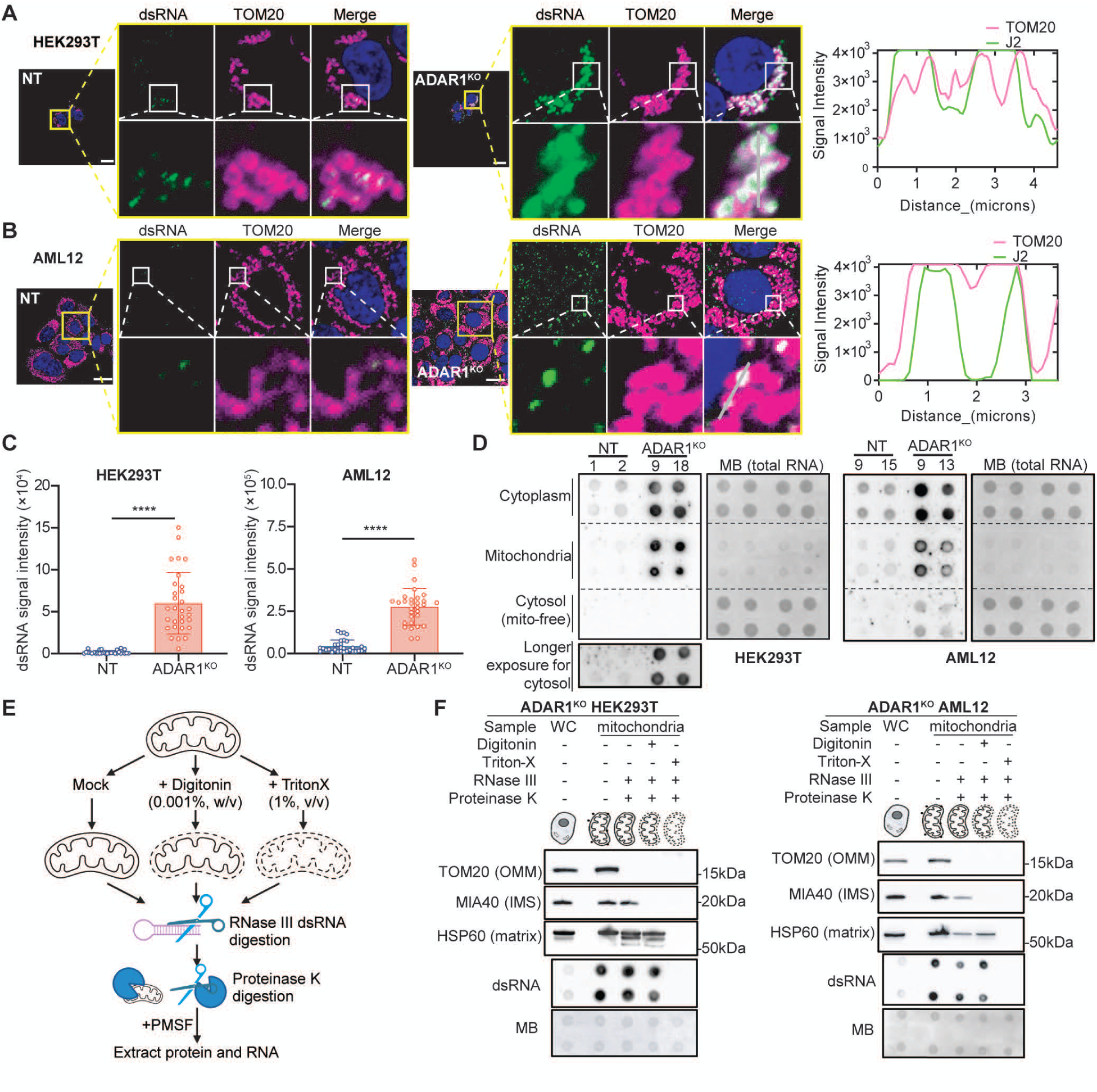
ADAR1 deficiency-induced dsRNAs accumulate in the mitochondrial matrix. (**A and B**) Representative immunofluorescence (IF) images showing mitochondria (TOM20, *magenta*) and dsRNA (J2, *green*) in NT and ADAR1^KO^ HEK293T (A) and AML12 (B) cells. Nuclei were counterstained by DAPI. Magnified views of the boxed regions are shown. Histograms on the right showing the fluorescence intensity along the grey lines drawn in the magnified views of (A) and (B), showing a high degree of coincidence between the signals corresponding to TOM20 and J2. Scale bars, 20μm. (**C**) Quantification of dsRNA signal intensity in NT and ADAR1^KO^ HEK293T (*left*) or AML12 (*right*) cells. Each dot represents an individual cell (*n* = 30 per group from three independent experiments). Data are presented as mean ± S.D. Statistical significance is determined using Mann-Whitney test (****, *P*<0.0001). (**D**) Dot blot analysis of dsRNA levels by J2 antibody in the indicated (sub)cellular fractions from NT and ADAR1^KO^ HEK293T (*left*) or AML12 (*right*) cells. MB staining was used as a loading control to verify the RNA sample input. The purity of the isolated cytoplasmic, cytosolic (mitochondria-free), and mitochondrial (mitochondria-enriched) fractions is shown in fig. S2F. (**E**) Workflow of the RNase III protection assay to determine dsRNA localization within mitochondria. (**F**) WB analysis of protein markers for each submitochondrial compartment, along with dot blot analysis of dsRNA levels, in the indicated samples from ADAR1^KO^ HEK293T (*left*) or AML12 (*right*). WC, whole cell; OMM, outer mitochondrial membrane; IMS, intermembrane space. (**E and F**) Schematics created in BioRender. Shen, H. (2025) https://BioRender.com/r25t501.

We next isolated cytoplasmic, cytosolic (mitochondria-free), and mitochondrial (mitochondria-enriched) fractions, followed by the dsRNA dot blot analysis in these fractions of both NT and ADAR1^KO^ HEK293T and AML12 cells (**fig. S2, E and F**). Similarly, we observed a striking enrichment of cellular dsRNAs in the mitochondrial fractions (**Fig. 2D**). The double-membrane structure of mitochondria defines three distinct compartments: the mitochondrial matrix, the intermembrane space (IMS), and the outer membrane surface (OMM). To further explore the localization and accessibility of dsRNAs induced by ADAR1 loss within different sub-mitochondrial compartments (**Fig. 2E**), we performed an RNase III protection assay using isolated mitochondria from ADAR1^KO^ HEK293T and ADAR1^KO^ AML12 cells under different membrane permeabilization conditions. Digitonin was used to selectively permeabilize the OMM (*28*), while Triton X-100 disrupted both the outer and inner membranes. The mitochondria were subsequently treated with RNase III and with proteinase K to further assess dsRNA accessibility (**Fig. 2, E and F**). Subsequent dsRNA dot blot and Western blot analyses showed that ADAR1 deficiency-induced dsRNAs were susceptible to RNase III digestion only when both mitochondrial membranes were permeabilized (**Fig. 2F**). These findings indicate that ADAR1 deficiency-induced mitochondria-localized dsRNA resides within the mitochondrial matrix.

### Mitochondrial matrix-localized dsRNAs, transcribed from the mitochondrial genome, contribute significantly to the endogenous dsRNA pool upon ADAR1 loss

It is known that mitochondria are a major source of self-dsRNA in mammalian cells, generated through bidirectional transcription from both strands of the mitochondrial circular genome, which frequently forms dsRNA intermolecularly (*29, 30*). Previous studies have suggested that ADAR1 suppresses cytosolic dsRNA sensing, primarily mediated by MDA5, with IR*Alu*s embedded in mRNAs transcribed from the nuclear genome being the main source of cytoplasmic dsRNAs (*8, 31*). However, our findings identify mitochondrial matrix-localized dsRNA as an additional major source of cellular dsRNAs following ADAR1 loss. We next investigated the origin of these mitochondrial matrix-localized dsRNA. To address this, we first treated ADAR1^KO^ cells with selective inhibitors targeting distinct mammalian RNA polymerases, including RNA polymerase I inhibitor *CX-5461* ^ref^ (*32*), RNA polymerase II/III inhibitor *α-amanitin* (*33*), RNA polymerase III inhibitor *ML-60218* ^ref^ (*34*), and mitochondrial RNA polymerase inhibitor *IMT-1* ^ref^ (*35*) (**Fig. 3, A and B**). Notably, treatment with *IMT-1*, but not other RNA polymerase inhibitors, substantially reduced the accumulation of mitochondrial matrix-localized dsRNA in ADAR1^KO^ cells, as shown by both dot blot analysis and immunofluorescence (IF) staining (**Fig. 3, A-C**). These results suggest that the mitochondrial matrix-localized dsRNAs accumulated in ADAR1^KO^ cells are transcribed from the mitochondrial genome, hence referred to as mitochondrial dsRNA (mt-dsRNA). To further unbiasedly investigate the origin of ADAR1 deficiency-induced cytoplasmic dsRNAs, we developed a dsRNA-eCLIP-seq method (double-stranded RNA enhanced cross-linking immunoprecipitation sequencing) to profile cytoplasmic dsRNAs in both NT and ADAR1^KO^ cells (**Fig. 3D**). We defined regions with significant enrichment in eCLIP samples compared to the size-matched input as “dsRNA peaks” (log_2_[Fold change _eCLIP vs Input_] > 1 and log_10_(*P*-value) < 2; see Methods). Within this set, regions with increased dsRNA peak density in ADAR1^KO^ cells compared to the NT counterparts were further classified as “upregulated dsRNA peaks” (log_2_[Fold change _KO vs NT_] > 1 and log_10_(*P*-value) < 2; see Methods). In HEK293T cells, this approach identified 1,728 dsRNA peaks, of which 1,122 are upregulated dsRNA peaks, confirming ADAR1’s role as a general dsRNA repressor (**fig. S3D** and **data S2**). Among these, 1,063 upregulated dsRNA peaks derived from the nuclear genome, with 52.63% and 18% being located in 3’UTR and introns, respectively (**fig. S3, B and C**). Further analysis showed that 89.87% of upregulated dsRNA peaks originated from short interspersed nuclear elements (SINEs), of which 99.58% were *Alu* elements (**fig. S3, C and D**). Intriguingly, similar to the nuclear genome (non-chrM)-aligned dsRNA reads, we observed a dramatic increase in total dsRNA reads aligned to the mitochondrial genome (chrM) in ADAR1^KO^ cells compared to NT counterparts (**fig. S3E**). Furthermore, we detected 61 dsRNA peaks on chrM, of which 59 were upregulated in ADAR1^KO^ compared to the NT cells (defined as “upregulated chrM peaks”) (**Fig. 3E**). Surprisingly, although chrM is a small circular genome (∼16.5 kb) compared to the nuclear genome, the chrM-aligned dsRNA reads showed an increased proportion of total dsRNA reads from approximately 65% in NT cells to 70% in ADAR1^KO^ cells (**Fig. 3F**). Moreover, the dsRNA peaks on chrM exhibited significantly higher fold changes in ADAR1^KO^ *versus* the NT control cells, relative to their non-chrM counterparts (**Fig. 3G**). Notably, the upregulated chrM dsRNA peaks derived from both the mitochondrial heavy and light strands, with a comparable distribution, suggesting that mt-dsRNAs likely form intermolecularly between transcripts from the two strands (**Fig. 3H**). To rule out cell line specificity, NT and ADAR1^KO^ AML12 cells were also subjected to the dsRNA-eCLIP-seq analysis, revealing similar changes (**Fig. 3, I-L, fig. S3, f and g,** and **data S2**). Altogether, our findings suggest that ADAR1 deficiency in both human and murine cells leads to accumulation of cytoplasmic dsRNAs derived not only from nuclear genome-transcribed transcripts but also from a substantial proportion of mt-dsRNAs, which predominantly form intermolecularly between the mitochondrial heavy and light strands.

**Fig. 3.**
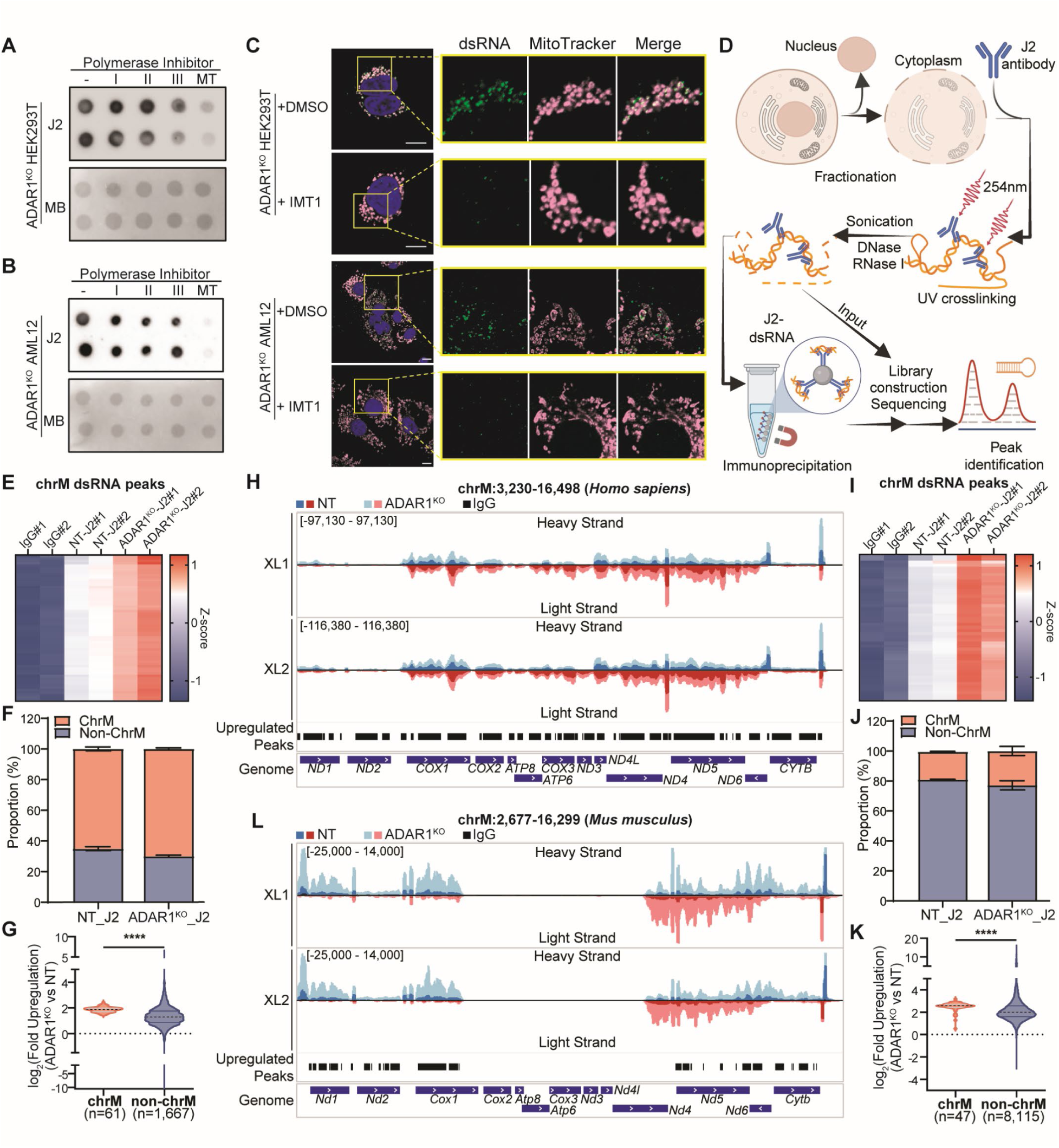
Mitochondrial matrix-localized dsRNAs, transcribed from the mitochondrial genome, contribute significantly to the endogenous dsRNA pool upon ADAR1 loss. (**A and B**) Dot blot analysis of dsRNA levels in isolated mitochondrial fractions from ADAR1^KO^ HEK293T (A) or AML12 (B) cells treated with the indicated RNA polymerase inhibitors. MB staining was used as a loading control. (**C**) Representative IF images showing dsRNAs (J2, *green*), mitochondria (Mitotracker, *magenta*) and nuclei (DAPI, *blue*) in ADAR1^KO^ HEK293T (*upper*) or AML12 (*lower*) cells treated with DMSO or IMT1. Magnified views of the boxed regions are shown. Scale bar, 10μm. (**D**) Schematic workflow of cytoplasmic dsRNA-eCLIP-seq. Two independent experiments were performed. Created in BioRender. Shen, H. (2025) https://BioRender.com/q61b383. (**E and I**) Heatmaps illustrating the difference in chrM dsRNA peak density across different eCLIP samples in HEK293T cells (E) and AML12 cells (I). Peak density data are row-normalized using z-scores for each peak, represented by a color spectrum, with each column representing the indicated eCLIP sample. (**F and J**) Proportions of chrM- and non-chrM dsRNA reads among total dsRNA reads detected in NT and ADAR1^KO^ HEK293T (F) or AML12 (J) cells. (**G and K**) Violin plot depicting fold changes in peak density for chrM and non-chrM dsRNA peaks in ADAR1^KO^ versus NT HEK293T (G) and AML12 (K) cells. The solid lines represent the quartiles, while the dashed lines indicate the median value. Statistical significance is measured by Mann-Whitney test (****, *P*<0.0001). (**H and L**) Integrated genome viewer (IGV) browser tracks of dsRNA reads mapped to chrM, from dsRNA-eCLIP-seq in HEK293T (H) and AML12 (L) cells. Data from two biological replicates (XL1 and XL2) are presented.

### ADAR1 represses mt-dsRNA through modulating mtROS level independent of its catalytic capability

ADAR1 is known to repress cellular dsRNA via its RNA binding and editing capabilities (*8, 23*). However, neither the nucleus-localized isoform p110 nor the nucleus- and cytoplasm-localized isoform p150 localizes to mitochondria (*36*), indicating a potential indirect regulation of ADAR1 on mt-dsRNA biogenesis. Notably, the increase in chrM dsRNA peaks following ADAR1 loss was not accompanied by the elevated expression of their host mitochondrial genes (**fig. S4A**). Additionally, no obvious expression change was observed in several previously reported regulators of mt-dsRNA, including SUV3 ^ref^ (*29*), PNPT1 ^ref^ (*29*), GRSF1 ^ref^ (*37*) and REXO2 ^ref^ (*38*) in ADAR1^KO^ cells (**fig. S4B**).

The accumulation of mt-dsRNA in the mitochondria of ADAR1^KO^ cells suggested a potential link to mitochondrial dysfunction. Mitochondrial function is tightly associated with oxidative phosphorylation and regulation of reactive oxygen species (ROS) (*39, 40*). Intriguingly, most of the mito-genes identified in the CRISPR screen are known to play established roles in ROS regulation. For instance, SOD2, OGDH, and SDHD directly control mtROS levels (*41–43*), while other mito-genes, such as SLC25A19, LIPT1, and SLC25A26, influence ROS indirectly by impacting various aspects of mitochondrial metabolism (*44–49*) (**Fig. 4A**). This association prompted us to investigate whether ADAR1 deficiency affects mtROS homeostasis in cells. We found that ADAR1^KO^ cells displayed significantly elevated mtROS level compared to the NT cells, indicating that loss of ADAR1 disrupts mitochondrial redox balance (**Fig. 4, B and C,** and **fig. S5, A and B**). To investigate whether manipulating mtROS could regulate mt-dsRNA, NT or ADAR1^KO^ cells were treated with either mitoParaquat (MitoPQ), a mitochondria-targeted redox cycler that directly and specifically induces mtROS (*50*), or MitoTEMPO, a mitochondria-targeted superoxide scavenger that specifically reduces mtROS (*51*), followed by measurement of mt-dsRNA levels. Strikingly, we observed a clear positive correlation between mtROS levels and mt-dsRNA levels in both HEK293T and AML12 cells (**Fig. 4, D-F, and fig. S5, C-G**). Moreover, treatment of the NT cells with 2 additional mtROS inducers - Antimycin A (a complex III inhibitor) and Oligomycin (a complex V inhibitor) similarly resulted in increased mt-dsRNA levels, providing further evidence for the role of mtROS in regulating mt-dsRNA accumulation (**fig. S5, D and E**). Our earlier data showed that silencing *SOD2*, one of the 6 validated mito-genes identified in our screen, triggered IFN signalling in ADAR1^KO^ cells (**Fig. 1, B and D**). Notably, these changes were mitigated by treatment with mitoTEMPO (**Fig. 4G**), which could effectively reduce mtROS levels in NT and ADAR1^KO^ cells (**fig. S5H**). These results suggest that mtROS plays a critical role in modulating dsRNA-induced innate immune activation in ADAR1-deficient cells.

**Fig. 4.**
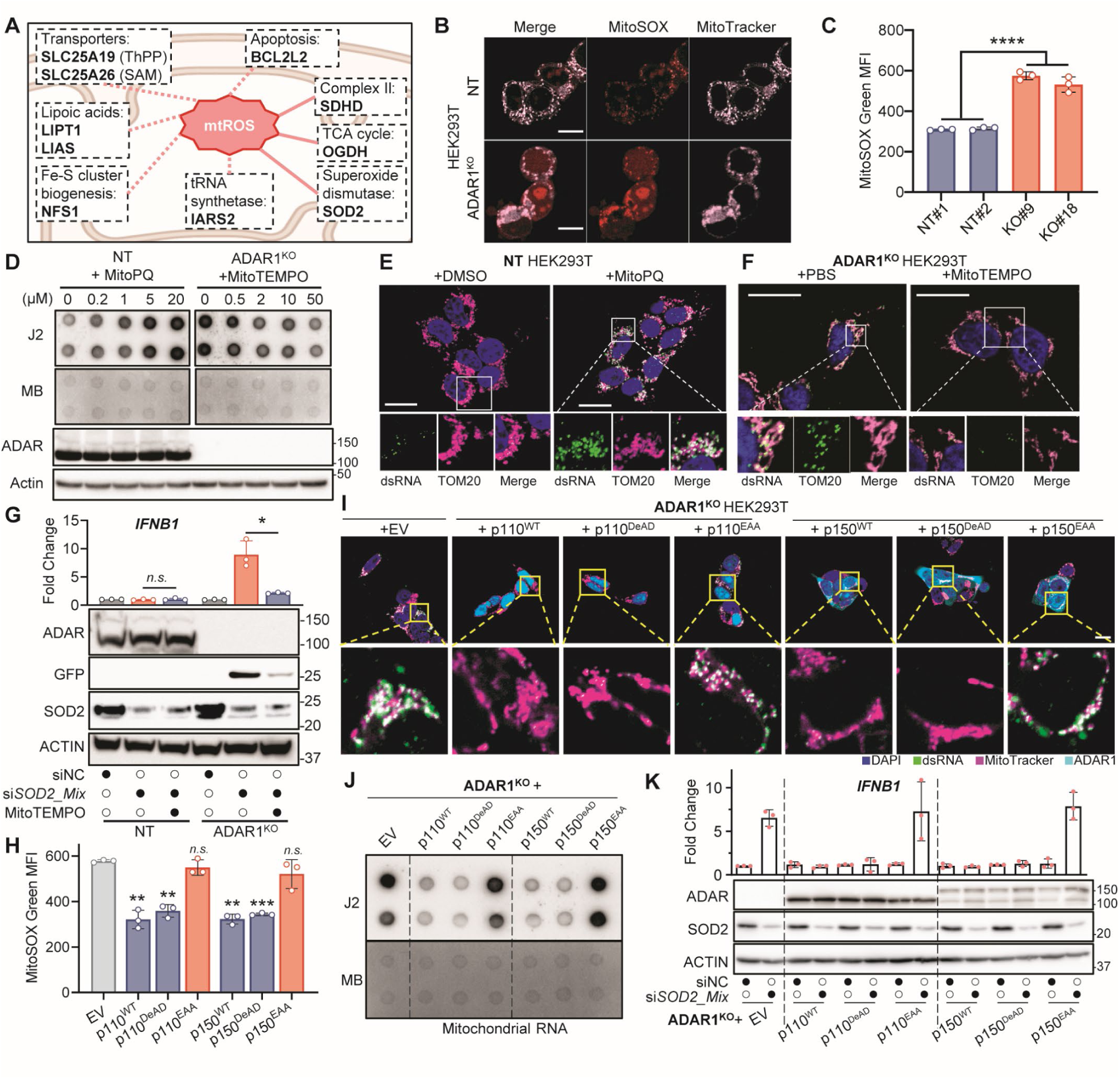
ADAR1 represses mt-dsRNA through modulating mtROS level independent of its catalytic capability. (**A**) Diagram summarizing the roles of identified mitochondrial immune protectors in the regulation of mitochondrial ROS (mtROS). Solid lines denote direct regulatory pathways, while dashed lines represent indirect mechanisms. Canonical protein functions are indicated above the respective gene names for clarity. (**B**) Representative images showing mtROS levels (MitoSOX-red, *red*) and mitochondria (MitoTracker, *magenta*) in NT and ADAR1^KO^ HEK293T cells. Scale bar, 10 μm. (**C**) FACS analysis of mtROS levels (stained by MitoSOX-green) in the indicated NT and ADAR1^KO^ HEK293T clones. MFI, median fluorescence intensity. Data are presented as the mean ± S.D. of 3 biological replicates. Statistical significance is measured by nested two-way ANOVA (****, *P*<0.0001). (**D**) Dot blot analysis of dsRNA levels in isolated mitochondrial fractions from the indicated cell groups, along with WB analysis of ADAR1 expression in protein lysates from the same cell groups. NT and ADAR1^KO^ HEK293T cells were treated with increasing concentrations of mitoPQ or mitoTEMPO for 16 hours, respectively. MB staining was used as a loading control. (**E and F**) Representative IF images showing dsRNAs (J2, *green*), mitochondria (TOM20, *magenta*) and nuclei (DAPI, *blue*) in NT cells treated with DMSO or mitoPQ (20 μM) for 16 hours (E), and in ADAR1^KO^ cells treated with PBS or mitoTEMPO (50 μM) for 16 hours (F). Magnified views of the boxed regions are shown. Scale bar, 20 μm. (**G**) QPCR analysis of *IFNB1* expression and WB analysis of GFP expression in NT and ADAR1^KO^ cells transfected with siNC or si*SOD2_Mix* (mixture of si*SOD2* #1 and si*SOD2*#2), with or without mitoTEMPO treatment (50 μM). (**H**) FACS analysis of mtROS levels (stained by MitoSOX-green) in ADAR1^KO^ HEK293T cells reintroduced with the indicated WT or mutant forms of ADAR1 isoforms. (**I**) Representative IF images showing dsRNAs (J2, g*reen*), mitochondria (Mitotracker, m*egenta*) and nuclei (DAPI, *blue*) in the same cell groups described in (H). Magnified views of the boxed regions are shown. Scale bar, 10 μm. (**J**) Dot blot analysis of dsRNA levels in isolated mitochondrial fractions from the same cell groups described in (H). MB staining was used as a loading control. (**K**) QPCR analysis of *IFNB1* expression and WB analysis of ADAR1 and SOD2 expression levels, in ADAR1^KO^ HEK293T cells reintroduced with the indicated WT or mutant forms of ADAR1 isoforms, along with transfection of siNC or si*SOD2_Mix.* (**G, H, and K**) Each dot represents the mean value of technical triplicates from an independent experiment. Data are presented as the mean ± S.D. of 3 biological replicates. Statistical significance is measured by paired, two-tailed Student’s *t*-test. *, *P*<0.05; **, *P*<0.01; ***, *P*<0.001; ****, *P*<0.0001; *n.s*., not significant, *P*>0.05.

Next, we investigated whether ADAR1’s known dsRNA-binding and editing functions (*52*) are essential for regulating mtROS levels and the resulting changes in mt-dsRNA levels within the cells. We overexpressed either the wild-type (WT) or mutant forms of ADAR1 p110 and p150 isoforms (DeAD, an editing-deficient mutant (*53*); EAA, a mutant lacking both RNA binding and editing abilities (*54, 55*)) in ADAR1^KO^ HEK293T cells. Subsequently, we measured mtROS and mt-dsRNA levels, as well as IFN signalling activation. Our results showed that, similar to the WT ADAR1 p110 and p150 isoforms, the DeAD mutant, but not the EAA mutant, was able to suppress mtROS and mt-dsRNA levels (**Fig. 4, H-J**), suggesting that ADAR1 reduces mtROS production and subsequent mt-dsRNA accumulation in an RNA binding-dependent but editing-independent manner. Furthermore, *IFNB1* upregulation triggered by *SOD2* depletion in ADAR1^KO^ cells was completely abolished by the restoration of either the WT or DeAD mutants, but not by the EAA mutants (**Fig. 4K**). These findings reveal an editing-independent functional axis of ADAR1 that prevents mtROS induction and mt-dsRNA accumulation, thereby averting aberrant IFN signalling and innate immune responses.

### ADAR1 deficiency-induced mt-dsRNA activates IFN signalling and innate immunity through a “Draw-and-Release” model

From the experiments described above, we observed that ADAR1 deficiency leads to elevated mtROS and subsequent accumulation of mt-dsRNA. However, as long as mitochondrial immune protectors identified in our screen remain intact, cells seem to tolerate these changes without activating dsRNA sensing or triggering IFN signalling and innate immune response. We therefore aimed to systematically investigate how mt-dsRNAs and mitochondrial proteins work in tandem to regulate the innate immune response following ADAR1 loss. This was achieved using multiple distinct treatment combinations to mimic different phases of the dynamic process, including: (1) KO of the *ADAR1* gene, (2) knockdown of *SOD2*, one of the validated mito-gene identified as a key protector against immune activation (**Fig. 1, B and D**), (3) treatment with MitoPQ, and (4) treatment with mitochondrial RNA polymerase inhibitor *IMT-1* (**Fig. 3, A and B**). Consistently, either ADAR1 KO or MitoPQ treatment alone led to a substantial accumulation of dsRNAs in mitochondrial but only a mild increase of dsRNA in cytosol, without causing any detectable immune activation (lanes #1, 2, 9, 10; **Fig. 5A**). However, with simultaneous knockdown of *SOD2*, we detected a significant increase in cytosolic dsRNA levels, accompanied by pronounced activation of IFN signalling, as indicated by elevated expression of *IFNB1* and ISGs (lanes #5, 6, 13, 14; **Fig. 5A**). Moreover, treatment of *IMT-1* suppressed both mitochondrial and elevated cytosolic dsRNA levels as well as the immune response under all tested conditions (lanes #7, 8, 15, 16; **Fig. 5A**). From these findings, it is very likely that defects in mitochondrial immune protectors may trigger the release of mt-dsRNAs to the cytosol. To test it, we examined the changes in subcellular distribution of mt-dsRNA in ADAR1^KO^ cells upon *SOD2* knockdown. In the siNC cells, cytoplasmic dsRNA signals were predominantly localized within mitochondria. However, upon *SOD2* knockdown, dsRNA signals were diffusely distributed into the cytosol (**Fig. 5B**). To specifically monitor the release of mt-dsRNAs into the cytosol triggered by SOD2 depletion, we performed fluorescence in situ hybridization (FISH) targeting the mitochondrial transcript *mt-Nd6* in ADAR1^KO^ AML12 cells. Following *Sod2* knockdown, we observed a significant increase in cytosolic *mt-Nd6* transcripts (**Fig. 5C**). Furthermore, RNA immunoprecipitation using J2 antibodies followed by quantitative PCR (J2 RIP-qPCR) revealed significant enrichment of chrM-encoded transcripts, but not the non-chrM-encoded transcripts *MOCOS* and *ZNF329*, in the cytosol of SOD2-depleted cells (**Fig. 5D**).

**Fig. 5.**
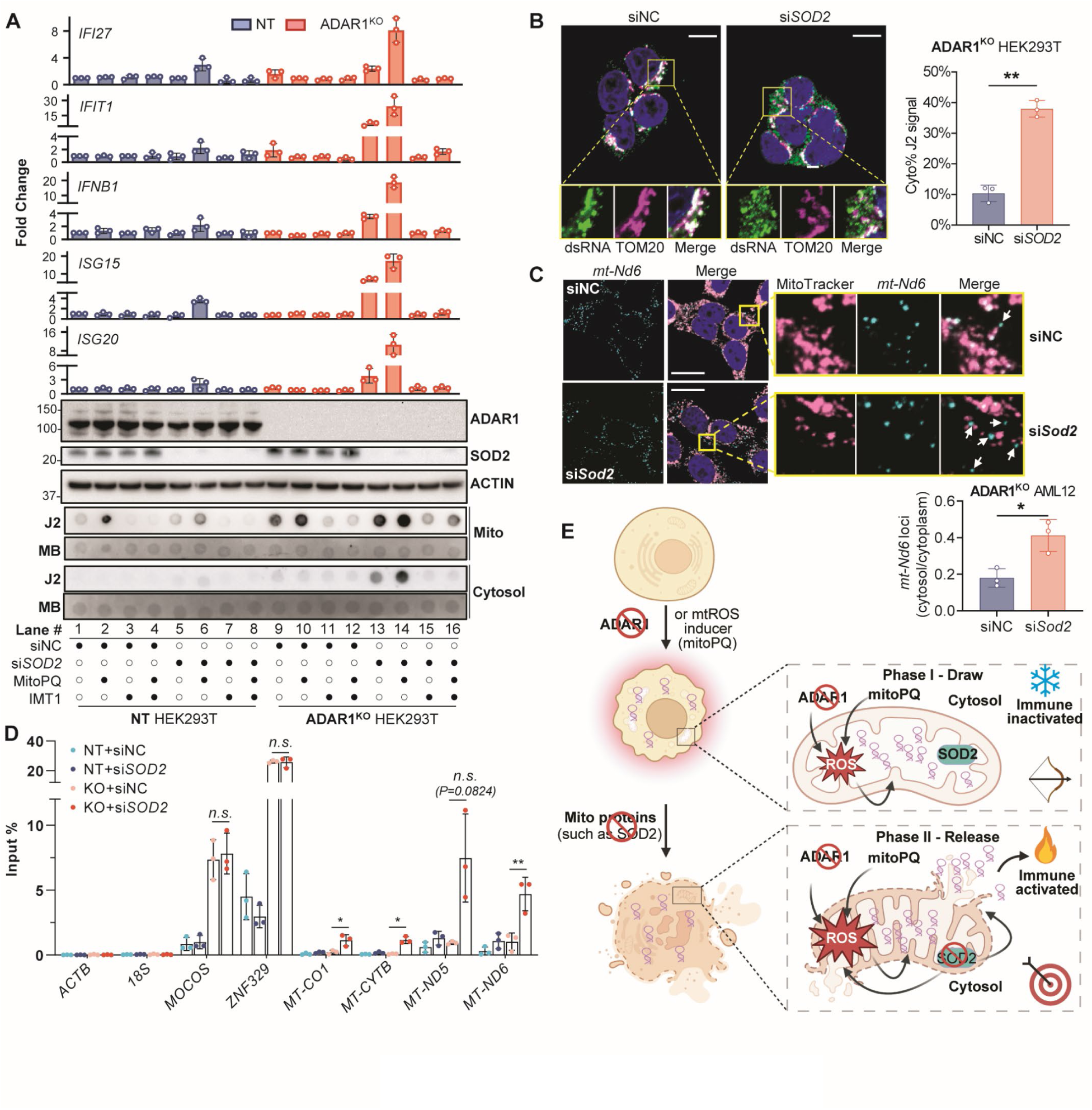
Mt-dsRNA induces RNA sensing and IFN response through a “*Draw*-and-*Release*” model. (**A**) QPCR analysis of the expression levels of *IFNB1* and *ISGs*, including *IFI27*, *IFIT1*, *ISG15*, and *ISG20* (*top*, bar charts) and WB analysis of ADAR1 and SOD2 expression (*middle*) using total RNA and protein samples from the indicated cell groups under different treatments. Dot blot analysis of dsRNA levels in the cytosolic and mitochondrial fractions of the indicated cell groups is shown in the bottom panel. For the bar charts, each dot represents the mean value of technical triplicates from an independent experiment. MB staining was used as a loading control. (**B**) Representative IF images showing dsRNAs (J2, *green*), mitochondria (TOM20, *magenta*) and nuclei (DAPI, *blue*) in ADAR1^KO^ HEK293T cells treated with siNC or si*SOD2*_mix. Bar charts showing the percentage of dsRNA signals detected in the cytosol (cytosol%) relative to the total dsRNA signals detected in the cytoplasm. Magnified views of the boxed regions are shown. Each dot represents the mean value per cell calculated from 4 fields of view. Scale bar, 10μm. (**C**) FISH images showing mitochondrial transcript *mt-Nd6* (*cyan*), mitochondria (Mitotracker, *magenta*) and nuclei (DAPI, *blue*) in ADAR1^KO^ AML12 cells upon treatment of siNC or si*Sod2*_mix (*top*). Bar charts (*bottom*-*right*) depicting the percentage of *mt-Nd6* foci formed in the cytosol relative to the total number of foci detected in the cytoplasm. Magnified views of the boxed regions are shown. Each dot represents the mean value per cell calculated from 5 fields of view. Scale bar, 20 μm. (**D**) RNA immunoprecipitation using J2 antibodies followed by qPCR (J2 RIP-qPCR) analysis of the expression levels of the indicated chrM- and non-chM-encoded transcripts in J2 pulldown products from the cytosolic fractions of NT or ADAR1^KO^ HEK293T cells treated with siNC or si*SOD2*_mix. Each dot represents the mean value of technical triplicates from an independent experiment. (**E**) Schematic illustration of the "*Draw*-and-*Release*" model. Created in BioRender. Shen, H. (2025) https://BioRender.com/m73b858. (**A to D**) Data are presented as the mean ± S.D. from 3 biological replicates. Statistical significance is determined using paired, two-tailed Student’s *t*-test. *, *P*<0.05; **, *P*<0.01; *n.s*., not significant.

All these findings led us to suggest a "*Draw*-and-*Release*" model describing this two-phase sequential process in which the absence of ADAR1 or the presence of mtROS-inducing agents triggers the occurrence of the "*Draw*" phase, characterized by elevated mtROS level that drives the accumulation of mt-dsRNA within the mitochondria. During this phase, mt-dsRNA remains confined to the mitochondrial matrix, preventing its interaction with cytosolic dsRNA sensors (**Fig. 5E**). However, when mitochondrial immune protectors, such as SOD2, are defective or downregulated, mitochondrial homeostasis is disrupted. This disruption leads to the release of mt-dsRNA into the cytosol, marking the “*Release*” phase. During this phase, cytosolic mt-dsRNA engages dsRNA-sensing pathways, culminating in immune activation (**Fig. 5E**).

### Reducing mtROS mitigates liver damage and inflammation in Adar1-deficient mice

Our data showed that following ADAR1 deficiency, elevated mtROS levels and the subsequent accumulation of mt-dsRNA contribute to dsRNA sensing and the IFN response only when mitochondrial gene depletion occurs. These responses can be suppressed by reducing mtROS, underscoring its pivotal role in driving innate immune activation in the absence of ADAR1. Moreover, we observed that mt-dsRNAs constitute a substantial portion of cellular dsRNAs induced by ADAR1 deficiency. This prompted us to investigate whether mtROS regulate mt-dsRNA level and if reducing mtROS could mitigate inflammation and pathologies associated with ADAR1 loss *in vivo*.

Our group recently reported that liver-specific deletion of the *Adar1* gene in mice (*Adar1*^flox/flox^ *Albumin*-Cre^+^) results in embryonic lethality, and concurrent deletion of one allele of the *Ifih1* gene (encoding MDA5) (*Ifih1*^+/-^*Adar1*^flox/flox^*Albumin-Cre*^+^) rescued this embryonic lethality. However, these mice still exhibited reduced body weight and shortened lifespans (*16*). In this study, we showed that *Ifih1*^+/-^*Adar1*^flox/flox^*Albumin-Cre*^+^ (referred to as *Ifih1*^+/-^*Adar*^f/f^) mice demonstrated severe hepatic inflammation and dysfunction, characterized by elevated blood aspartate transaminase (AST) and alanine transaminase (ALT) levels, disrupted liver hepatic architecture, ballooning hepatocytes, and prominent inflammatory infiltrates, especially CD68- and CD8-positive cells (**fig. S6, A-D**). In addition, the liver is particularly vulnerable to both ADAR1 deficiency and mitochondrial stress (*56, 57*). Our earlier data from murine hepatocyte AML12 cells support this vulnerability by demonstrating elevated mtROS and subsequent mt-dsRNA accumulation following ADAR1 deficiency. Together, these observations highlight *Ifih1*^+/-^ *Adar*^f/f^ mice as an optimal *in vivo* model to validate the “*Draw*-and-*Release*” process. To this end, we first isolated primary hepatocytes from perfused livers of *Ifih1*^+/-^*Adar*^wildtype/wildtype^*Albumin-Cre*^+^ (referred to as *Ifih1*^+/-^*Adar*^wt/wt^) and *Ifih1*^+/-^*Adar*^f/f^ mice. Our findings revealed that primary hepatocytes from *Ifih1*^+/-^*Adar*^f/f^ mice exhibited significantly higher levels of mtROS elevation, dsRNA accumulation in both cytosol and mitochondria, and ISG upregulation, compared to the *Ifih1*^+/-^*Adar*^wt/wt^ counterparts (**Fig. 6, A-C, fig. S6E,** and **data S3**). Further analysis revealed overall reduced expression levels of identified mitochondrial immune protectors in hepatocytes from *Ifih1*^+/-^*Adar*^f/f^ mice (**Fig. 6D**). This finding underscores the potential involvement of mitochondrial dysfunction in the pathology of these mice and suggests the possible release of mt-dsRNAs into the cytosol.

**Fig. 6.**
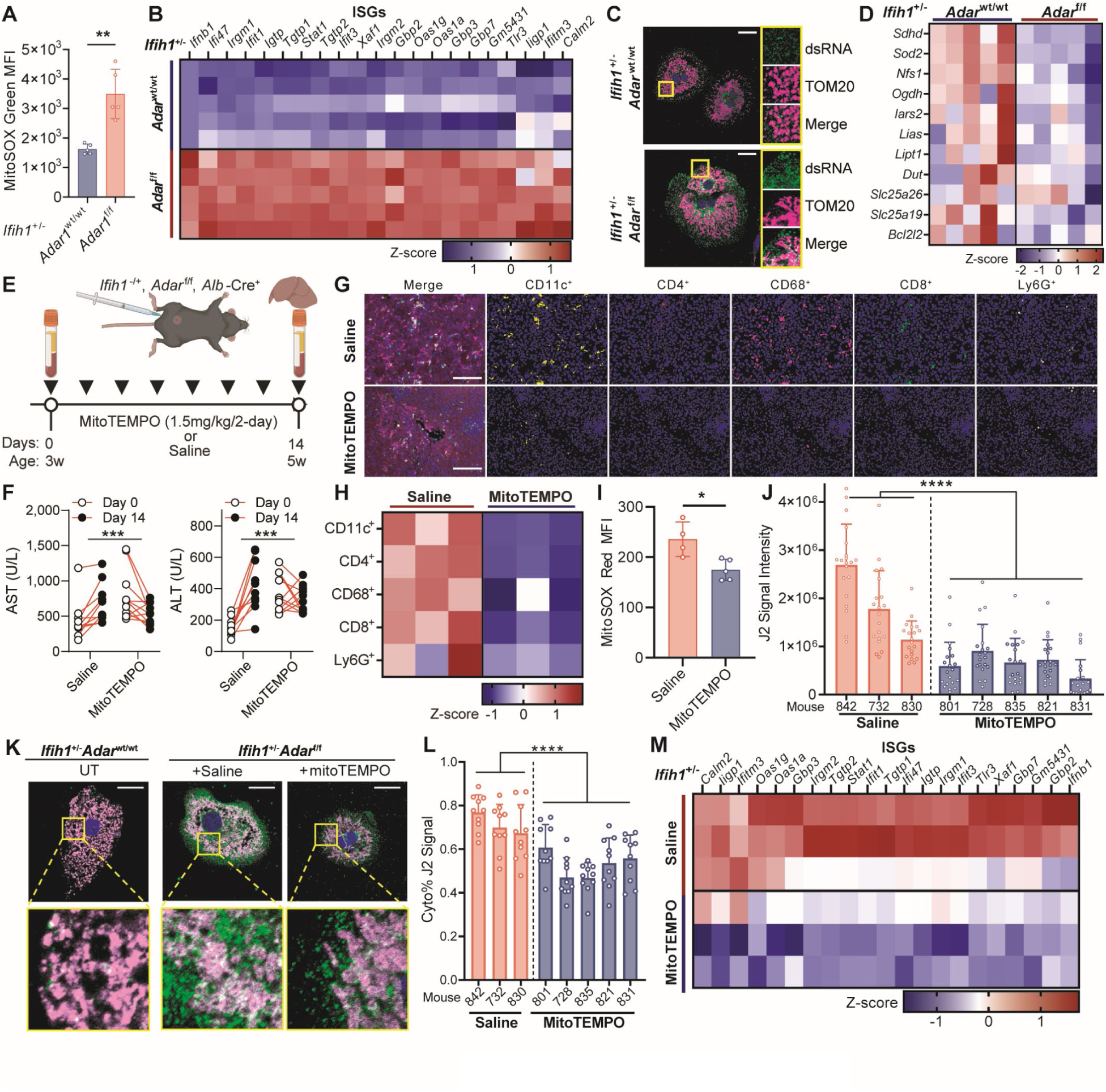
Treatment reducing mtROS alleviated hepatic damage following ADAR1 ablation in mice. (**A**) FACS analysis of mtROS levels (stained by MitoSOX-green) in primary hepatocytes from the indicated mouse groups (*n* = 5 mice per genotype). (**B**) Heatmap illustrating the expression levels of ISGs based on RNA-seq analysis of primary hepatocytes from the indicated mouse groups (*n* = 5 mice per genotype). Expression data are column-normalized using z-scores for each gene, represented by a color spectrum, with each row representing an individual mouse. (**C**) Representative IF images showing dsRNA (J2, *green*), mitochondria (TOM20, *magenta*) and nuclei (DAPI, *blue*) in primary hepatocytes from the indicated mouse groups. Magnified views of the boxed regions are shown. Scale bar, 20 μm. (**D**) Heatmap showing the expression levels of mito-genes identified from our screen in the same samples described in (B). (**E**) Schematic diagram of the mitoTEMPO treatment experiment in mice. Created in BioRender. Shen, H. (2025) https://BioRender.com/v68b856. (**F**) Liver damage marker tracing of aspartate aminotransferase (AST) (*left*) and alanine aminotransferase (ALT) (*right*) in the saline-treated (n = 10) and mitoTEMPO-treated (n = 11) mouse groups before and after treatment (Day 0 and Day 14). Statistical significance is determined by two-way repeated measures ANOVA. (**G**) Representative multiplexed fluorescent IHC (mfIHC) images of liver tissue infiltrates. Merged images show pseudo-colored staining for CD11c (*yellow*), CD4 (*red*), CD68 (*magenta*), CD8 (*green*), Ly6G (*pink*), and DAPI (*blue*), labeled as “Merge.” Additional images display cell segmentation masks and marker positivity (indicated at the top), with blue representing negative staining and other colors indicating positive staining. Scale bars, 100 μm. (**H**) The composition profiles of the tissue infiltrates in the liver of from the mouse with indicated treatment (*n* = 3 for each group) as determined by mfIHC. The proportions were row-normalized using z-score, represented by a color spectrum, with each column representing an individual mouse. (**I**) FACS analysis of mtROS levels (stained by MitoSOX-Red) in primary hepatocytes from the indicated treatment groups (*n* = 4 or 5 for the ‘saline’ or ‘mitoTEMPO’ group, respectively). (**J**) Quantification of dsRNA signal intensity in primary hepatocytes from mice in the ‘saline’ or ‘mitoTEMPO’ group (*n*=3 or 5, respectively). Each dot represents an individual cell (*n* = 20 per mouse). (**K**) Representative IF images showing dsRNA (J2, *green*), mitochondria (TOM20, *magenta*) and nuclei (DAPI, *blue*) in primary hepatocytes isolated from one representative mouse in the ‘untreated’ (UT), ‘saline’ or ‘mitoTEMPO’ group. Magnified views of the boxed regions are shown. Scale bar, 20 μm. (**L**) Bar charts showing the percentage of dsRNA signals detected in the cytosol (cytosol%) relative to the total dsRNA signals detected in the cytoplasm of primary hepatocytes from the same mice described in (J). Each dot represents an individual cell (*n* = 10 per mouse). (**M**) Heatmap showing the expression changes of ISGs upon mitoTEMPO treatment based on RNA-seq analysis of primary hepatocytes from the indicated mouse groups (*n*=3 mice per group). Expression data are column-normalized using z-scores for each gene, represented by a color spectrum, with each row representing an individual mouse. (**A, I, J, and L**) Data are presented as mean ± S.D. Statistical significance is determined using two-tailed Welch’s *t*-test (A), two-tailed Student’s *t*-test (I), or nested two-way ANOVA (J and L). *, *P*<0.05; **, *P*<0.01; ***, *P*<0.001; ****, *P*<0.0001.

Next, diseased *Ifih1*^+/-^*Adar*^f/f^ mice received intraperitoneal administration of mitoTEMPO (1.5 mg/kg every 2 days) or the saline control (**Fig. 6E**). Phenotypically, in mice receiving the saline control, both AST and ALT levels increased over 14 days, whereas MitoTEMPO-treated mice exhibited stable or even reduced levels over the same period (**Fig. 6F**). Moreover, histological analysis revealed attenuated inflammation and hepatocyte damage following mitoTEMPO treatment, characterized by significant decrease of all tested inflammatory infiltrates, and improved hepatic architecture (**Fig. 6, G and H, and fig. S6F**). Primary hepatocytes isolated from mitoTEMPO-treated mice demonstrated reduced mtROS levels and dsRNA accumulation compared to the control counterparts (**Fig. 6, I and J**). Further analysis of the subcellular localization of dsRNAs in primary *Ifih1*^+/-^*Adar*^f/f^ hepatocytes revealed that mitoTEMPO treatment not only significantly reduced total dsRNA levels but also decreased the proportion of cytosolic dsRNAs, leading to a reduction in ISG expression (**Fig. 6, K-M, fig. S6G,** and **data S4**). These findings validate the "*Draw*-and-*Release*" model *in vivo*.

## Discussion

ADAR1 has emerged as a vital immune protector, orchestrating cytosolic innate immunity and safeguarding against inappropriate immune activation. A key question in the field is identifying the primary sources of immunogenic dsRNAs and uncovering the mechanisms governing cellular dsRNA sensing, particularly in the context of ADAR1 deficiency and its pathological consequences. Our study unveiled a more intricate and nuanced landscape, challenging conventional paradigms. We discovered that in both human and murine cells, ADAR1 deficiency induces accumulation of cellular dsRNAs derived not only from the well-characterized nuclear genome-transcribed IR*Alu*s embedded in 3′UTRs (*13, 58*) but also from a substantial proportion of mt-dsRNAs transcribed from the mitochondrial genome. Our study highlights a dual role for ADAR1 in maintaining cellular immune homeostasis. Beyond its well-established function as the “last line of defense” in cytosolic innate immunity, preventing unnecessary and aberrant dsRNA sensing, ADAR1 also suppresses the production and accumulation of mt-dsRNA, thereby mitigating the risk of its release into the cytosol under cellular stress conditions.

Until now, the relationship between ADAR1, mtROS, and mt-dsRNA has been largely overlooked, given that ADAR1 is not a mitochondria-localized protein (*36*). We showed that loss of ADAR1 results in elevated mtROS levels and the subsequent accumulation of mt-dsRNAs in the mitochondrial matrix, independent of ADAR1’s RNA editing capability. Notably, both the nucleus-localized ADAR1p110 and the predominantly cytoplasm-localized ADAR1p150 isoforms require their dsRNA-binding activity to rescue the mtROS elevation caused by ADAR1 deficiency to a similar extent. This suggests that regardless of its subcellular localization, ADAR1 likely functions as an RNA-binding protein (RBP) rather than as an editing enzyme to indirectly regulate mtROS levels through interactions with other RBPs that are known to regulate mtROS, such as human antigen R (HuR) (*59, 60*) and Y-box binding protein 1 (YB1) (*61*).

mtROS not only act as signalling molecules that can induce oxidative conditions but also directly oxidize biomolecules, including RNA, which is more susceptible to oxidative damage than DNA (*62*). Oxidized mitochondrial RNA has been proved to be able to promote inflammatory responses in human THP-1 monocyte cells (*^63^*), although detailed mechanisms remain unknown. Our study demonstrated that mtROS elevation induces the formation of mt-dsRNA, highlighting the need for further investigation into the mechanistic link between mtROS-induced RNA oxidation and the structural changes in mitochondrial RNA that facilitate intermolecular mt-dsRNA formation. Notably, oxidative stress, as reported in yeast, can cause RNA-protein cross-linking (*64*), potentially impairing mitochondrial RNA degradation and contributing to mt-dsRNA accumulation. Proteins like ADAR1 may play a regulatory role in managing oxidized RNA levels, balancing their immunogenic potential and functional roles as part of an adaptive cellular mechanism.

Despite the well-established role of mitochondria in immune regulation, the mechanisms by which mt-dsRNAs become accessible to cytosolic RNA sensors to drive immune responses remains largely unexplored, particularly in the context of ADAR1 deficiency—a condition linked to autoimmune disorders (*20*). A recent study reported that ADAR1 deficiency in hepatocellular carcinoma (HCC) cells disrupts mitochondria structure and function, characterized by shrinkage, increased membrane density, reduced cristae, and rupture of the mitochondrial outer membrane (*65*). Here, we propose a “*Draw*-and-*Release*” model: during the "*Draw*" phase, elevated mt-dsRNA levels are sequestered within mitochondria, and during the "*Release*" phase, these dsRNAs are released into the cytosol upon depletion of mitochondrial immune protectors, triggering immune responses. This mechanism, supported by both *in vitro* cell line studies and *in vivo* mouse models, highlights a conserved and generalizable role for ADAR1 in regulating mt-dsRNA levels across species. Interestingly, mitochondria have retained distinct prokaryotic features such as their circular genome and bacterial-type RNA synthesis machinery, enabling the formation of intermolecular mt-dsRNA, which can serve as potent RNA ligands for cellular immune sensors (*2, 66, 67*). Cytosolic presence of mt-dsRNAs has been implicated in triggering IFN production by activating RIG-I and MDA5 (*29*). A recent study suggested that cytosolic RNA can directly activate MAVS by interacting with its intrinsically disordered domain (*68*). Given that MAVS is anchored on OMM, it is also plausible that released mt-dsRNA could directly induce an IFN response through MAVS.

Therapeutically, treatment with the mtROS scavenger MitoTEMPO significantly reduced hepatic immune activation and tissue damage in liver-specific ADAR1 KO mice, highlighting the potential of mtROS modulation as a therapeutic strategy for ADAR1-associated pathologies, including Aicardi-Goutières syndrome and other interferonopathies. Intriguingly, our findings also demonstrate that even in the presence of functionally intact ADAR1 protein, chemical-mediated mtROS levels increase coupled with depletion of mitochondrial proteins such as SOD2 is capable of triggering an innate immune response. This immune activation is driven by the release of accumulated mt-dsRNA into the cytosol, likely overwhelming ADAR1’s capacity to surveil dsRNA. This discovery reveals an alternative pathway for activating immune responses, particularly in cancer cells. By concurrently inducing mtROS and promoting the release of mt-dsRNAs through inhibiting mitochondrial proteins such as SOD2, it may be possible to circumvent ADAR1-mediated suppression of cytoplasmic dsRNA sensing, presenting a novel strategy for anti-cancer immunotherapies.

## Acknowledgments

We thank and acknowledge the Microscopy and Multiplex Assays (MMA) Core Facility and the Fluorescence Activated Cell Sorting (FACS) Facility at the Cancer Science Institute of Singapore for their assistance with imaging and FACS-based experiments. This work was supported by National Research Foundation Singapore; Singapore Ministry of Education under its Research Centres of Excellence initiative.

## Author contributions

Conceptualization: L.C., H.S.

Methodology: H.S., J.H., V.T., E.T.

Investigation: H.S., V.T., J.H., V.H.E.N., W.L.G., S.Q.P.

Visualization: H.S., V.T.

Funding acquisition: L.C.

Resources: J.H., S.J.T., W.L.G., L.N., B.Y.L.N., K.W.L., X.S., J.Z.

Formal analysis: H.S., V.T.

Supervision: L.C.

Writing – original draft: H.S., L.C., V.T.

Writing – review & editing: H.S., L.C.

## Supplementary Materials

Materials and Methods

Figs. S1 to S6

References (*69–79*)

Data S1 to S5

## Materials and Methods

### Ethical statement

Our research complies with all relevant ethical regulations. All animal experiments were approved by the Institutional Animal Care and Use Committees of National University of Singapore (NUS; Singapore) with the protocol numbers BR23-0476 and R23-0457.

### Mice strains

*Adar*^flox^ mutant mice (B6.129-*Adar* ^tm1Knk^/Mmjax) and *Ifih1*^−/−^ mice (B6.Cg-*Ifih1*^tm1.1Cln^/J) were kindly provided by the laboratory of Associate Professor Sun Lei. Alb-Cre mice (B6.Cg-Speer6-ps1^Tg(Alb-cre)21Mgn^/J) were purchased from the Jackson laboratory and used for breeding. For genotyping, ear tips were collected from mice, and genomic DNA was extracted using DirectPCR (Ear) lysis reagent (Viagen #402-E) with 1% Proteinase K (Thermo Fisher Scientific AM2546) following the manufacturer’s protocol. PCR was carried out using Platinum Green 2X PCR Master Mix (Invitrogen 13001012). Genotyping was performed according to the protocols provided by The Jackson Laboratory. All mice used in this study were of C57BL/6J background, and comparisons were made using age- and gender-matched controls. All mice were maintained in pathogen–free (SPF) facility in NUS Comparative Medicine Department. Less than 5 mice with same sex were housed in a cage at 20-25 °C and 50% humidity with a 12-hour light/dark cycle.

### Mouse blood liver biomarker measurement

Submandibular blood collection was performed on *Ifih1^+/−^Adar^wt/wt, flox/flox^Alb-Cre^+^* mice at indicated time points. Whole blood was collected in MiniCollect Complete CAT Serum Separator Clot Activator tubes (Greiner, 450548). The samples were then submitted to the Comparative Medicine Diagnostic Lab at NUS for analysis of liver biomarkers, including AST and ALT.

### Mouse intraperitoneal administration of mitoTEMPO

*Ifih1^+/−^Adar ^flox/flox^Alb-Cre^+^*mice were randomly assigned to two groups, receiving either treatment of MitoTEMPO (Merck, SML0737) at a dose of 1.5 mg/kg MitoTEMPO (Merck, SML0737) in sterile saline, or same volume of sterile saline control, via intraperitoneal (i.p.) injection every two days for 14 days. Submandibular blood collection was performed both before the first injection and after the final injection, as previously described. On day 14, at the end of the experiment, mice were sacrificed for downstream analyses.

### Two-Step Collagenase Liver Perfusion and Hepatocyte Isolation

Mouse livers were perfused using a two-step collagenase digestion. Following perfusion with prewarmed perfusion buffer (1× HBSS; Gibco, 14180046) containing 0.5 mM EGTA (pH 8; 1^st^ Base, CUS-1070) and digestion buffer (serum-free DMEM containing 0.05% collagenase 1A; Sigma-Aldrich, C9891), the softened liver was excised, dissected, and hepatocytes were released, filtered, and centrifuged. Red blood cells were lysed using Ack lysing buffer (Gibco, A10492-01), and live hepatocytes were isolated using a 10% Percoll (Sigma-Aldrich, P1644) centrifugation at 200 × g. Cells were washed with with cold DMEM containing 10% FBS and 0.5 U/mL RQ1 RNase-free DNaseI (Promega, M6101), resuspended in FBS-supplemented DMEM, and prepared for experiments.

### Primary Mouse Hepatocyte Culture

Primary hepatocytes isolated via two-step collagenase liver perfusion were seeded onto glass coverslips in 6-well plates at a density of 1 × 10⁵ cells per well, or 6-well plates at a density of 7 × 10⁵ cells per well. Plates and glass coverslips were pre-coated with 1% Matrigel (Merck, CLS356231) in serum-free DMEM/F-12 medium for 30 minutes at 37°C. Hepatocytes were cultured in DMEM/F-12 supplemented with 10% FBS.

### Cell lines

EC109 cells (RRID:CVCL_6898) were kindly provided by Professor TSAO, George Sai Wah (Director, Faculty Core Facility, Li Ka Shing Faculty of Medicine); HEK293T (Cat#ATCC-CRL-3216; RRID:CVCL_0063), AML12 (Cat#ATCC-CRL-2254; RRID:CVCL_0140), and Hepa1-6 (Cat#CRL-1830; RRID:CVCL_0327) cells were purchased from the American Type Culture Collection (ATCC). HEK293T cells and Hepa1-6 were maintained in Dulbecco’s Modified Eagle Medium (Biowest); AML12 cells were maintained in Dulbecco’s Modified Eagle Medium/Nutrient Mixture F-12 (Gibco); EC109 cells were maintained in Roswell Park Memorial Institute (RPMI) medium (Biowest), all supplemented with 10% fetal bovine serum (FBS; Thermo Scientific) and incubated at 37°C with 5% CO_2_. Drugs including MitoTEMPO (Merck, SML0737), MitoPQ (Abcam, ab146819), Antimycin A (Merck, A8674), Oligomycin (MedChemExpress, HY-N6782), CX-5461 (MedChemExpress, HY-13323), ML-60218 (MedChemExpress, HY-122122), α-Amanitin (MedChemExpress, HY-19610), and IMT1 (MedChemExpress, HY-134539) were added into cell medium with the concentration and duration as described.

### RNA extraction and RT-qPCR

RNA from cells were extracted using RNeasy mini kit (Qiagen) with on column treatment of DNaseI, or FastPure Cell/Tissue Total RNA Isolation Kit V2 (Vazyme, RC112). cDNA was synthesized using Advantage RT-for-PCR kit (Clontech) with random hexamer primers and subsequently qPCR was performed with GoTaq DNA polymerase (Promega). Fold change is calculated by 2^-ΔΔCt_sample_. ΔCt= Ct_target_ – Ct_actin;_ ΔΔCt=ΔCt_sample_-averageΔCt_control_. Primers used in RT-qPCR are listed in Data S5.

### RNA sequencing and gene expression analysis

RNA samples were extracted from isolated hepatocytes samples and sent to BGI for DNBSEQ Eukaryotic Long Non-Coding RNA sequencing. Genome alignment and read counting of the RNA-sequencing data was performed using RSEM (*69*) with options for STAR (v2.7.4a) (*70*) paired end alignment to the Ensembl mm9 reference genome assembly and gene annotation gtf (NCBIM37 release 67 *Mus musculus*). For differential gene expression analysis, the gene count matrix was imported into R and analysed using the DESeq2 R package (*71*) (v1.42.1, R v4.3.3). Briefly, gene expression analysis was performed using the DESeq function with options for Wald test. Where available, sex is included as a covariate in the linear model design to account for any confounding sex effects. DEGs were filtered using the following cutoffs: mean counts≥20, |logFC|≥1, and adj. *P*-value<0.05. Gene Ontology functional term Gene Set Enrichment Analysis was performed using the clusterProfiler R package (v4.10.1) (*72*).

### Western blot

Cells were lysed in RIPA buffer (Sigma, R0278) supplemented with 1× cOmplete EDTA-free protease inhibitor cocktail (Roche, 11873580001). Total protein concentrations were determined using the Bradford assay (Bio-Rad, 5000006). Proteins were separated on 10% SDS-PAGE gels and transferred onto polyvinylidene difluoride (PVDF) membranes (Millipore, IPVH00010). Membranes were incubated with primary antibodies (1:1,000 dilution) overnight at 4 °C, followed by incubation with secondary antibodies (1:10,000 dilution) for 1 hour at room temperature. Blots were visualized using enhanced chemiluminescence (GE Healthcare, RPN2106). Antibodies used in this study are as listed: anti-ADAR1 (CST, #14175; for detecting human ADAR1), anti-ADAR1 15.8.6 (Santa Cruz, sc-73408; for detecting mouse ADAR1), anti-β actin HRP (Santa Cruz Biotechnology, sc-47778HRP), anti-mouse IgG HRP-linked (Cell Signaling Technology, 7076), anti-rabbit IgG HRP-linked (Cell Signaling Technology, 7074), anti-GFP (Santa Cruz, SC-9996), anti-MDA5 (Cell Signaling Technology, 5321), anti-TBK1 (Cell Signaling Technology, 3504), anti-MAVS (Cell Signaling Technology, 3993), anti-IRF3 (Cell Signaling Technology, 11904), anti-NFκB (Cell Signaling Technology, 8242T), anti-TOM20 (ProteinTech, 11802-1-AP), anti-SUV3 (ProteinTech, 12826-1-AP), anti-PNPT1 (ProteinTech, 14487-1-AP), anti-HSP60 (ProteinTech, 66041-1-Ig), anti-MIA40 (ProteinTech, 21090-1-AP), anti-SOD2 (ProteinTech, 66474-1-Ig), anti-GRSF1 (abcam, ab205531), anti-REXO2 (Abcam, ab206694), anti-Alpha-Tubulin (Santa Cruz, sc-32293), anti-Fibrillarin (Abcam, ab4566).

### Transfection and lentivirus production

Transfection was carried out with Lipofectamine 2000 (Thermo Fisher Scientific) according to the manufacturer’s protocol, at a ratio of 3μL per 1μg plasmids. For lentivirus generation, HEK293FT cells in a T75 flask were co-transfected with 5μg viral vectors, 2.5μg of pRSV-Rev, 2.5μg of pMD2.G, and 7.5μg pMDLg/pRRE. The medium constaining lenti-virus was then harvested 48 hours after transfection. Medium was then filtered followed by concentration using Lenti-X Concentrator (Takara Bio) and resuspended with DMEM and stored at -80℃.

Dicer-substrate RNAs (DsiRNAs) were ordered from Integrated DNA Technologies (IDT). A total of 25nM of siRNAs were transfected into HEK293T or AML12 cells with Lipofectamine RNAiMax (Thermo Fisher Scientific) at a ratio 1μL per 0.01nmol siRNA. The cells were collected by scraping 48 h later for downstream experiments. Sequences of siRNAs are provided in Data S5.

### Generation of ADAR1 knockout (KO) cells

The NT and *ADAR1*^KO^ HEK293T cell lines were established using lipofectamine-mediated transfection with lentiCRIPSR v2 (Addgene) plasmid containing non-targeting gRNA sequences or gRNA sequences targeting human *ADAR1*. The NT or *Adar1*^KO^ EC109, Hepa1-6, and AML12 cell lines were established using lentiviral transduction with plentiCRIPSR v2 (Addgene) plasmid containing non-targeting gRNA sequences or gRNA sequences targeting mouse *Adar1*, followed by puromycin selection. Single clones of transfected or transduced cells were used for further experiments. All clones were validated with sequencing showing genome editing and western blotting showing depletion of ADAR1 protein. Guide RNA sequences are provided in Data S5.

### Generation of *IFNB1*-reporter cells

The 1,500bp promoter region of human *IFNB1* gene was amplified from human placenta genomic DNA (Sigma) using Q5 HS HiFi DNA polymerase (NEB, M0493L) while GFP DNA together with Kozak sequence was amplified from pEGFP-N1 plasmid (Clontech) using Q5 HS HiFi DNA polymerase. The two DNA were then ligated together and cloned into a pLenti6/V5-DEST plasmid (Invitrogen, V49610) replacing the original CMV-enhancer, CMV-promoter, and V5-tag. The *IFNB1*-reporter was then packed into lentivirus and transduced into HEK293T cells at multiplicity of infection (MOI) = 0.1, followed Blasticidin S selection and single clone isolation.

### CRISPR Screening

Human GeCKOv2 CRISPR knockout pooled library was a gift from Feng Zhang (Addgene #1000000048). CRISPR screening was carried out as described (*73*). A total of 120 × 10^6^ NT or ADAR1^KO^ HEK293T cells were infected with lentivirus carrying library A at MOI = 0.3, followed by puromycin (1μg/ml) selection, resulting in coverage >500. Infected cells were then cultured for one week, and expended to 100 × 10^6^ cells for FACS sorting on BD FACSAria II Cell Sorter. Sorting was performed based on GFP signal (FITC channel). Cells with the top 2% and the bottom 5% GFP signal were sorted and collected as “IFN-high” population and “IFN-low” population, respectively. After sorting, the cells from both populations were subjected for genomic DNA extraction using the Zymo Quick-DNA™ Miniprep Plus Kit (Zymo Research, D4068) according to the manufacturer’s protocol. The integrated sgRNAs were amplified (first PCR) and Illumina adapters were added (second PCR) using Q5 HS HiFi DNA polymerase (NEB, M0493L). The PCR products were purified and sequenced using HiSeq X by Novogene. The enrichment of screen candidate genes was analysed using MAGeCK (*57*) by comparing enrichment of sgRNA in “IFN-high” population versus “IFN-low” population. The analysis was performed on usegalaxy.org (*74*), following the tutorial provided. Significantly enriched hits were defined as log2FoldChange >1 and FDR<0.05. Significantly enriched hits in ADAR1^KO^ cells were input to g:Profiler (version e110_eg57_p18_4b54a898) for GO analysis.

### Cell fractionation and mitochondria isolation

For Extended Figure 1b and cytoplasmic J2-eCLIP sequencing, cell fractionations were performed using PARIS™ Kit (Invitrogen) with the manufacturer’s instrument.

For other relevant figures except Figure 5a, cell fractionation followed a Dounce homogenization method (*75*). Briefly, cell pellets were resuspended in 1.1 mL ice-cold RSB hypo buffer (10 mM NaCl, 1.5 mM MgCl₂, 10 mM Tris-HCl, pH 7.5) and and allowed to swell for 5–10 minutes before lysis with a B pestle. The homogenate was diluted with 0.8 mL 2.5× MS buffer (525 mM mannitol, 175 mM sucrose, 12.5 mM Tris-HCl, 2.5 mM EDTA, pH 7.5) and subjected to differential centrifugation. Nuclear and cellular debris were removed by centrifugation at 700g for 5 minutes, repeated twice. 10% of the supernatant were saved as cytoplasmic fraction. The remaining of the supernatant was centrifuged at 13,000g for 15 minutes, with pellet as crude mitochondria fraction, and supernatant as cytosol fraction. Mitochondria were washed with 1× MS buffer three times, while cytosol were centrifuged three times. All solutions, tubes, and homogenizers were prechilled on ice, and all centrifugation steps were conducted at 4°C.

For Figure 5a, a rapid approach suitable for isolating mitochondria from multiple samples was modified from a previous study (*28*). Briefly, cells from a 6cm dish were harvested by trypsinization, counted, and washed with PBS. After pelleting, cells were resuspended in ice-cold RSB hypo buffer with 0.02% digitonin (Merck, D141) and incubated on ice for 5 min while vortexing every minute for 5 min. A 2/3 volume of 2.5× MS buffer was added, and the samples were centrifuged at 700g for 5 min to remove debris, repeated twice. The supernatant was then centrifuged at 13,000g for 15 min, with pellet as crude mitochondria fraction, and supernatant as cytosol fraction. Mitochondria were washed with 1× MS buffer three times, while cytosol were centrifuged three times.

### RNase III digestion

For dot blot analysis, 10µg RNA samples were digested with 5U RNase III (Invitrogen, AM2290) for 30 minutes at 37°C according to manufacturer’s protocol, followed by RNA extraction using RNeasy mini kit (Qiagen).

For immunofluorescence analysis, after permeabilization (see “Immunofluorescence” section below), 0.5U RNase III was added into PBS with 10 mM MgCl_2_ and incubated for 30 minutes at 37°C, followed by washing three times with PBS.

For both analyses, parental samples were subjected to the same processing without addition of RNase III (undigested).

### RNase III Protection Assay

Freshly isolated mitochondria from cells were resuspended in 1x MS homogenization buffer and divided into four tubes (A, B, C, D) for differential treatments. Tube C was permeabilized with 0.001% digitonin, while others added the same volume of 1x MS buffer. Following incubation on ice and centrifugation, mitochondria were resuspended in 1x MS buffer with 5 mM MgCl₂, with tube-specific additions: no RNase III for tube A, RNase III (10 U/mL) for tubes B and C, and RNase III plus 1% Triton-X100 for tube D. Samples were incubated at 37°C for 15 minutes, followed by treatment of Superase-In and proteinase K, and supplemented with BME and PMSF before centrifugation. Pellets were washed, transferred, and processed for RNA and protein extraction, with tube D handled differently due to complete lysis by Triton-X100.

### Immunofluorescence

Cells were cultured on glass coverslips at a density of 10% 24 hours before. For experiments using MitoTracker, MitoTracker™ Deep Red FM (Invitrogen) was added in growth media to a final concentration of 500 nM and incubating for 30 min at 37°C prior to fixation. Cells were then fixed in 4% paraformaldehyde in PBS for 15 minutes at room temperature, and permeabilized with 0.2% Triton X-100 in PBS for 10 minutes. After blocking with 5% bovine serum albumin (BSA) in PBS for 30 minutes, samples were incubated with primary antibodies (dilution 1:200) overnight at 4°C. Coverslips were washed with PBS and incubated with secondary antibodies (dilution 1:200) for 1 hour at room temperature in the dark. Coverslips were mounted with VECTASHIELD® Antifade Mounting Medium with DAPI (Eon Biotech, H-1200-10). Fluorescence images were acquired using a Nikon A1R confocal microscope with magnification at 60× or 100×. Fluorescence intensities were analysed using FIJI. Primary antibodies used: anti-dsRNA J2 primary antibody (Nordic Mubio, 10010500), anti-ADAR1 (CST, #14175), anti-TOM20 (Proteintech, 11802-1-AP). Secondary antibodies used: goat anti-Rabbit secondary antibody, Rhodamine Red™-X (Invitrogen, R-6394) and goat anti-Mouse secondary antibody, Alexa Fluor 488 (Abcam, ab150113).

### Dot Blot assays

For dsRNA detection, 2 µL of RNA samples per dot was applied to Amersham Hybond-NX nylon membrane (Cytiva). Each dot contains 50 ng for RNA extracted from mitochondria, 1 µg for RNA extracted from whole cell, cytoplasm, or cytosol. The samples were then crosslinked with 120 mJ/cm² of UV light (254 nm). After blocking with 5% non-fat dry milk in PBST for 30 minutes, the membranes were incubated with the anti-dsRNA J2 primary antibody (dilution 1:1,000) overnight at 4°C. Following washes with PBST, the membranes were incubated with secondary antibody (dilution 1:10,000) for 1 hour at room temperature. Enhanced chemiluminescence (GE Healthcare) was used to visualize the blots.

### mtROS Measurement

For fluorescence imaging, cells were seeded in a µ-Slide 8 well high chamber (Ibidi) and incubated with MitoSOX Red (5 µM; Invitrogen) and MitoTracker™ Deep Red FM (500 nM; Invitrogen) in medium at 37°C for 30 minutes. Fluorescence images were acquired using a Nikon A1R confocal microscope.

For flow cytometry, cells were stained with MitoSOX Green (500 nM; Invitrogen) or MitoSOX Red (5 µM; Invitrogen) in medium at 37°C for 30 minutes. Cells were washed twice with PBS, resuspended in 1x PBS, and analysed immediately using a FACSymphony™ A3 Cell Analyzer (BD) with FITC channel. Data analysis was performed using FlowJo™ v10 (BD). Gating strategy for flow cytometry analysis was provided in Figure S1.

### Hematoxylin and Eosin (H&E) Staining

H&E staining was performed on formalin-fixed paraffin-embedded (FFPE) liver tissue sections. Tissues were fixed in 4% paraformaldehyde in PBS overnight at room temperature and embedded in paraffin. Sections were cut at 4 μm thickness and deparaffinized with Clearene (Leica, 3803600) through three washes. Rehydration was achieved by sequential immersion in ethanol solutions of decreasing concentration (2 × 100%, 90%, 80%, and 70%) for 5 minutes each. Slides were stained with hematoxylin (Leica, 3801522) for 1 minute, rinsed in running water for 1 minute, treated with acid-alcohol for 1 minute, rinsed again, and stained with eosin (Leica, 3801601) for 1 minute. After washing in running water, sections were dehydrated using ethanol solutions of increasing concentration (70%, 80%, 90%, and 2 × 100%), followed by two washes in Clearene. Slides were mounted with CV Ultra mounting medium (Leica, 14070936261), coverslipped, and imaged using the Olympus BX43 imaging system.

### Automated mfIHC Staining and Quantification

Formalin-fixed paraffin-embedded (FFPE) tissue slides were processed using the Opal Multiplex fIHC kit (Akoya Biosciences, California) by the Microscopy and Multiplex Assays Core (CSI, NUS) with an automated protocol for CD4/CD8/CD68/CD11C/Ly6G staining was performed using the Leica Bond III (SN: 3214062) with the Bond Polymer Refine Detection Kit (Cat No: DS9800). Staining parameters (e.g., antibody concentration, incubation time, antigen retrieval pH/time) were optimized using traditional DAB staining. For multiplex fluorescence staining, slides were baked, dewaxed, and subjected to heat-induced epitope retrieval (HIER) at 100°C using HIER 1 or HIER 2 solution. After 10 min peroxidase blocking (only for the 1st marker), markers prepared in antibody diluent (Agilent, Cat No: DKO.S302283) were sequentially applied along with polymeric HRP-conjugated secondary antibody (LeicaBiosystems, DS9800) and opal fluorophores (Opal 480–620 from Akoya Bioscience; CF®680R from Biotium). Slides were rinsed with wash buffer between steps. After each marker, slides were heated at 100°C to strip bound antibodies for subsequent marker labelling.

Antibody-Opal Pairs and Conditions:

- CD4 (Abcam, ab183685, 1:500, HIER 1, 20 min) – Opal 480
- CD68 (Abcam, ab125212, 1:1000, HIER 1, 20 min) – Opal 520
- CD11C (CST, 97585, 1:100, HIER 2, 20 min) – Opal 570
- Ly6G (BD Pharmingen, 551459, 1:50, HIER 2, 20 min) – Opal 620
- CD8 (CST, 9849s, 1:200, HIER 2, 20 min) – CF®680R

For Ly6G (rat antibody), goat anti-rat IgG (HRP) was used for signal amplification. The Leica Bond III processed two antibodies per cycle, with reagents loaded between cycles. DAPI (Spectral DAPI, Akoya, FP1490, 1:10) was added as a nuclear counterstain, and slides were mounted in ProLong Diamond Antifade Mountant (P36961) and cured at room temperature for 24 h. Slides were imaged using the PhenoImager HT (Akoya Biosciences, S/N: VP2314N2212), and signal unmixing, cell segmentation, and phenotyping were conducted with inForm Software (version 2.6.0, Akoya Biosciences). Multiplex fluorescence images were analyzed with Visiopharm. DAPI nuclear staining were used to segment cells. The percentage of positively stained cells for each marker was quantified using the score function in inForm Software. For each image, cells that have indicated marker intensity above a given intensity threshold for that image were declared to be positive for that marker using inForm Software. The raw percentages are normalized using z-scores (*z* = (*x*-*μ*)/*σ*), where *z* is the normalized value (z-score), *x* is the raw percentage for a given immune cell marker in a sample, *μ* is the mean raw percentage across all samples for that marker, and *σ* is the standard deviation of the raw percentages across all samples for that marker.

### Cytoplasmic dsRNA-eCLIP sequencing

Cytoplasmic dsRNA-eCLIP sequencing was modified from standard eCLIP sequencing (*76*). Cytoplasm of the same number of cells from a 15 cm dish was fractionated using 500 μL PARIS cell fraction buffer, topped up to 1.5mL with lysis buffer (50 mM Tris pH 7.4, 100 mM NaCl, 1% NP40, 0.1% SDS, 0.5% sodium deoxycholate, protease inhibitor) and precleared with Dynabeads™ Protein G (Invitrogen, 10004D) coated with mouse IgG (Santa Cruz Biotechnology, sc-2025). Samples were incubated with 5 μg J2 dsRNA antibody or mouse IgG, UV crosslinked (400 mJ/cm², 254 nm), sonicated (Bioruptor on the low setting in 4°C for 5 min, cycling 30 seconds on and 30 seconds off), and treated with TURBO™ DNase (Invitrogen, AM2238) and Ambion™ RNase I (Invitrogen, AM2295). RNA-protein complexes were immunoprecipitated using Dynabeads™ Protein G, washed with high salt buffer (50 mM Tris pH 7.4, 1 M NaCl, 1 mM EDTA, 1% NP40, 0.1% SDS, 0.5% sodium deoxycholate) and wash buffer (20 mM Tris pH 7.4, 10 mM MgCl2, 0.2% Tween-20), and digested with Proteinase K. RNA was extracted using TRIzol™ (Invitrogen, 15596018) and Zymo RNA Clean & Concentrator Kit (Zymo Research, R1019), dephosphorylated, adapter-ligated, reverse-transcribed, and sequenced on BGI DNBseq-G400. Two independent eCLIP were performed for each sample.

eCLIP-Seq read files were pre-processed and analysed using the eCLIP-seq CLIPper peak enrichment pipeline (*77*) with the to hg19 (Gencode GRCh37) and mm9 (Gencode NCBIM37). Peak enrichment scores for each CLIP samples were normalised using the corresponding input samples. Enriched peaks were defined as peaks with log_2_(Fold change) > 1 and log_10_(*P*-value) < 2. Functional and repeat element annotation of enriched peaks in the HEK293T and AML12 was performed using HOMER annotatePeaks using the hg19 and mm9 genome annotation, respectively (*78*). For differential peak analysis, enriched peaks from all samples were concatenated and merged for read depth analysis using samtools bedcov (v1.5) and imported into R for subsequent analyses. To account for potential systematic enrichment in high complexity genomic regions and library size effects, read counts of CLIP samples were subtracted by counts in the corresponding input samples at each peak and normalised using library size factors calculated from the input samples (*79*). Differential binding analysis was performed using the DESeq2 R package (v1.42.1) (*71*). The statistical significance threshold for differential binding was set at absolute log_2_Fold change > 1 and BH-adjusted *P*-value < 0.01.

### RNA immunoprecipitation (RIP)

Cytosolic fraction from same number of cells from a 10 cm dish were collected and 1% cytosolic fraction was saved as input. Cytosolic fractions were precleared with Dynabeads™ Protein G (Invitrogen) coated with normal mouse IgG (Santa Cruz Biotechnology, sc-2025) for 2 hours at 4°C with addition of 1% murine RNase inhibitor (NEB, M0314), followed by incubation with 2.5μg J2 dsRNA antibody or normal mouse IgG for 3 hours at 4°C. A total of 60ul Dynabeads™ Protein G was added into each sample and rotate in 4°C overnight. After immunoprecipitation, beads were washed with high salt wash buffer (50mM Tris pH7.4, 1M NaCal, 1mM EDTA, 1% NP40, 0.1% SDS, 0.5% sodium deoxycholate) twice and wash buffer (20mM Tris pH7.4, 10mM MgCl2, 0.2% Tween-20) twice. Input and IP samples were then digested with 20 μL Proteinase K in 130 μL Proteinase K buffer (100mM Tris pH7.4, 50mM NaCl, 10mM EDTA, 0.2% SDS) at 37°C 30 minutes, 50°C 30 minutes, 1,200 rpm. RNA was extracted with TRIzol™ (Invitrogen, 15596018) and Zymo RNA Clean & Concentrator Kits (Zymo Research, R1019). Extracted RNA was reverse-transcribed using Advantage RT-for PCR kit (Clontech) with random hexamer followed by qPCR. %Input = 2^-ΔCt^ ÷ Dilution Factor ×100%; ΔCt= Ct_RIP_ – Ct_input_. Sequences of primers are listed in Data S5.

### RNA fluorescence in situ hybridization

Cells were seeded into a µ-Slide 8 well high chamber (Ibidi) and RNA fluorescence in situ hybridization (FISH) was performed using the ViewRNA ISH Assay Kit (Thermo Fisher Scientific) following the manufacturer’s protocol. ViewRNA mouse ND6 Alexa Fluor 488 probes (ThermoFisher Scientific, VB4-3116021-PF) targeting mouse *mt-Nd6* was used. Nuclei were counterstained with DAPI (1 µg/mL), and fluorescence images were acquired using a Nikon A1R confocal microscope with magnification at 100×. *mt-Nd6* focis were analysed using FIJI (v1.54f).

### Statistical analyses

For two-sample comparisons, data for each sample were first tested for normality using the Shapiro-Wilk test. If one of the samples did not pass the test (*P* < 0.05 in Shapiro-Wilk test), the Mann-Whitney test was applied. If both samples followed a normal distribution (*P* > 0.05 in Shapiro-Wilk test), a two-tailed Student’s *t*-test was used when the F-test indicated equal variances (*P* > 0.05 in F-test). If the F-test showed significantly different variances (*P* < 0.05 in F-test), a two-tailed Welch’s *t*-test was applied. A nested two-way ANOVA was performed using the *aov* function in R (v4.3.3), while other statistical tests were conducted using GraphPad Prism (v10.4.1), unless otherwise specified.

## References and Notes

1. K. B. Chiappinelli et al., Inhibiting DNA methylation causes an interferon response in cancer via dsRNA including endogenous retroviruses. Cell 162, 974–986 (2015).

2. A. Dhir et al., Mitochondrial double-stranded RNA triggers antiviral signalling in humans. Nature 560, 238–242 (2018).

3. E. A. Bowling et al., Spliceosome-targeted therapies trigger an antiviral immune response in triple-negative breast cancer. Cell 184, 384–403 e321 (2021).

4. B. J. Liddicoat et al., RNA editing by ADAR1 prevents MDA5 sensing of endogenous dsRNA as nonself. Science 349, 1115–1120 (2015).

5. H. Chung et al., Human ADAR1 prevents endogenous RNA from triggering translational shutdown. Cell 172, 811–824. e814 (2018).

6. Y. Gao et al., m6A modification prevents formation of endogenous double-stranded RNAs and deleterious innate immune responses during hematopoietic development. Immunity 52, 1007–1021.e1008 (2020).

7. C. R. Capshew, K. L. Dusenbury, H. A. Hundley, Inverted Alu dsRNA structures do not affect localization but can alter translation efficiency of human mRNAs independent of RNA editing. Nucleic Acids Res 40, 8637–8645 (2012).

8. B. J. Liddicoat et al., RNA editing by ADAR1 prevents MDA5 sensing of endogenous dsRNA as nonself. Science 349, 1115–1120 (2015).

9. N. M. Mannion et al., The RNA-editing enzyme ADAR1 controls innate immune responses to RNA. Cell Rep 9, 1482–1494 (2014).

10. E. Eisenberg, E. Y. Levanon, A-to-I RNA editing - immune protector and transcriptome diversifier. Nat Rev Genet 19, 473–490 (2018).

11. L. Chen, A-to-I editing prevents self-RNA sensing. Nat Rev Mol Cell Biol 24, 85 (2023).

12. G. I. Rice et al., Mutations in ADAR1 cause Aicardi-Goutieres syndrome associated with a type I interferon signature. Nat Genet 44, 1243–1248 (2012).

13. P. Mehdipour et al., Epigenetic therapy induces transcription of inverted SINEs and ADAR1 dependency. Nature 588, 169–173 (2020).

14. C. M. McEntee, A. N. Cavalier, T. J. LaRocca, ADAR1 suppression causes interferon signaling and transposable element transcript accumulation in human astrocytes. Front Mol Neurosci 16, 1263369 (2023).

15. T. Zhang et al., ADAR1 masks the cancer immunotherapeutic promise of ZBP1-driven necroptosis. Nature 606, 594–602 (2022).

16. W. L. Gan et al., Hepatocyte-macrophage crosstalk via the PGRN-EGFR axis modulates ADAR1-mediated immunity in the liver. Cell Rep 43, 114400 (2024).

17. J. B. Saenz, N. Vargas, C. J. Cho, J. C. Mills, Regulation of the double-stranded RNA response through ADAR1 licenses metaplastic reprogramming in gastric epithelium. JCI Insight 7, (2022).

18. J. I. Kim et al., RNA editing at a limited number of sites is sufficient to prevent MDA5 activation in the mouse brain. PLoS Genet 17, e1009516 (2021).

19. T. Sun et al., A Small Subset of Cytosolic dsRNAs Must Be Edited by ADAR1 to Evade MDA5-Mediated Autoimmunity. bioRxiv, 2022.2008.2029.505707 (2022).

20. Q. Li et al., RNA editing underlies genetic risk of common inflammatory diseases. Nature 608, 569–577 (2022).

21. O. Solomon et al., RNA editing by ADAR1 leads to context-dependent transcriptome-wide changes in RNA secondary structure. Nat Commun 8, 1440 (2017).

22. J. E. Heraud-Farlow, et al., GGNBP2 regulates MDA5 sensing triggered by self double-stranded RNA following loss of ADAR1 editing. Sci Immunol 9, eadk0412 (2024).

23. S. B. Hu et al., ADAR1p150 prevents MDA5 and PKR activation via distinct mechanisms to avert fatal autoinflammation. Mol Cell 83, 3869–3884 e3867 (2023).

24. W. Li et al., MAGeCK enables robust identification of essential genes from genome-scale CRISPR/Cas9 knockout screens. Genome Biol 15, 554 (2014).

25. M. Zhang et al., TRAF-interacting protein (TRIP) negatively regulates IFN-beta production and antiviral response by promoting proteasomal degradation of TANK-binding kinase 1. J Exp Med 209, 1703–1711 (2012).

26. C. Uggenti et al., cGAS-mediated induction of type I interferon due to inborn errors of histone pre-mRNA processing. Nat Genet 52, 1364–1372 (2020).

27. A. Lepelley et al., Mutations in COPA lead to abnormal trafficking of STING to the Golgi and interferon signaling. J Exp Med 217, (2020).

28. B. Dixit, S. Vanhoozer, N. A. Anti, M. S. O’Connor, A. Boominathan, Rapid enrichment of mitochondria from mammalian cell cultures using digitonin. MethodsX 8, 101197 (2021).

29. A. Dhir et al., Mitochondrial double-stranded RNA triggers antiviral signalling in humans. Nature 560, 238–242 (2018).

30. Y. G. Chen, S. Hur, Cellular origins of dsRNA, their recognition and consequences. Nat Rev Mol Cell Biol 23, 286–301 (2022).

31. H. Chung et al., Human ADAR1 Prevents Endogenous RNA from Triggering Translational Shutdown. Cell 172, 811–824 e814 (2018).

32. M. J. Bywater et al., Inhibition of RNA polymerase I as a therapeutic strategy to promote cancer-specific activation of p53. Cancer Cell 22, 51–65 (2012).

33. M. Noe Gonzalez, D. Blears, J. Q. Svejstrup, Causes and consequences of RNA polymerase II stalling during transcript elongation. Nat Rev Mol Cell Biol 22, 3–21 (2021).

34. C. R. K, R. Cheng, S. Zhou, S. Lizarazo, D. J. Smith, K. Van Bortle, Evidence of RNA polymerase III recruitment and transcription at protein-coding gene promoters. Mol Cell, (2024).

35. A. Hooftman et al., Macrophage fumarate hydratase restrains mtRNA-mediated interferon production. Nature 615, 490–498 (2023).

36. S. Rath et al., MitoCarta3.0: an updated mitochondrial proteome now with sub-organelle localization and pathway annotations. Nucleic Acids Res 49, D1541–D1547 (2021).

37. F. Hensen et al., Mitochondrial RNA granules are critically dependent on mtDNA replication factors Twinkle and mtSSB. Nucleic Acids Res 47, 3680–3698 (2019).

38. M. Szewczyk et al., Human REXO2 controls short mitochondrial RNAs generated by mtRNA processing and decay machinery to prevent accumulation of double-stranded RNA. Nucleic Acids Res 48, 5572–5590 (2020).

39. M. T. Lin, M. F. Beal, Mitochondrial dysfunction and oxidative stress in neurodegenerative diseases. Nature 443, 787–795 (2006).

40. I. Martinez-Reyes, N. S. Chandel, Mitochondrial TCA cycle metabolites control physiology and disease. Nat Commun 11, 102 (2020).

41. R. J. Mailloux, D. Craig Ayre, S. L. Christian, Induction of mitochondrial reactive oxygen species production by GSH mediated S-glutathionylation of 2-oxoglutarate dehydrogenase. Redox Biol 8, 285–297 (2016).

42. S. Drose, Differential effects of complex II on mitochondrial ROS production and their relation to cardioprotective pre- and postconditioning. Biochim Biophys Acta 1827, 578–587 (2013).

43. J. A. Bolduc, J. A. Collins, R. F. Loeser, Reactive oxygen species, aging and articular cartilage homeostasis. Free Radic Biol Med 132, 73–82 (2019).

44. S. L. Ong, H. Vohra, Y. Zhang, M. Sutton, J. A. Whitworth, The effect of alpha-lipoic acid on mitochondrial superoxide and glucocorticoid-induced hypertension. Oxid Med Cell Longev 2013, 517045 (2013).

45. Z. Jiang, SLC25A19 is required for NADH homeostasis and mitochondrial respiration. Free Radic Biol Med 222, 317–330 (2024).

46. M. L. Hartman, M. Czyz, BCL-w: apoptotic and non-apoptotic role in health and disease. Cell Death Dis 11, 260 (2020).

47. Q. Dong et al., IARS2 mutations lead to Leigh syndrome with a combined oxidative phosphorylation deficiency. Orphanet J Rare Dis 19, 305 (2024).

48. G. P. Cheng, S. M. Guo, Y. Yin, Y. Y. Li, X. He, L. Q. Zhou, Aberrant Expression of Mitochondrial SAM Transporter SLC25A26 Impairs Oocyte Maturation and Early Development in Mice. Oxid Med Cell Longev 2022, 1681623 (2022).

49. S. W. Alvarez et al., NFS1 undergoes positive selection in lung tumours and protects cells from ferroptosis. Nature 551, 639–643 (2017).

50. M. P. Murphy et al., Guidelines for measuring reactive oxygen species and oxidative damage in cells and in vivo. Nat Metab 4, 651–662 (2022).

51. R. Ni et al., Therapeutic inhibition of mitochondrial reactive oxygen species with mito-TEMPO reduces diabetic cardiomyopathy. Free Radic Biol Med 90, 12–23 (2016).

52. K. Licht, M. F. Jantsch, The Other Face of an Editor: ADAR1 Functions in Editing-Independent Ways. Bioessays 39, (2017).

53. F. Lai, R. Drakas, K. Nishikura, Mutagenic analysis of double-stranded RNA adenosine deaminase, a candidate enzyme for RNA editing of glutamate-gated ion channel transcripts. J Biol Chem 270, 17098–17105 (1995).

54. L. Valente, K. Nishikura, RNA binding-independent dimerization of adenosine deaminases acting on RNA and dominant negative effects of nonfunctional subunits on dimer functions. J Biol Chem 282, 16054–16061 (2007).

55. L. Qi et al., An RNA editing/dsRNA binding-independent gene regulatory mechanism of ADARs and its clinical implication in cancer. Nucleic Acids Res 45, 10436–10451 (2017).

56. J. C. Hartner, C. Schmittwolf, A. Kispert, A. M. Muller, M. Higuchi, P. H. Seeburg, Liver disintegration in the mouse embryo caused by deficiency in the RNA-editing enzyme ADAR1. J Biol Chem 279, 4894–4902 (2004).

57. P. Middleton, N. Vergis, Mitochondrial dysfunction and liver disease: role, relevance, and potential for therapeutic modulation. Therap Adv Gastroenterol 14, 17562848211031394 (2021).

58. S. Ahmad et al., Breaching Self-Tolerance to Alu Duplex RNA Underlies MDA5-Mediated Inflammation. Cell 172, 797–810 e713 (2018).

59. I. X. Wang, E. So, J. L. Devlin, Y. Zhao, M. Wu, V. G. Cheung, ADAR regulates RNA editing, transcript stability, and gene expression. Cell Rep 5, 849–860 (2013).

60. J. H. Lee, M. Jung, J. Hong, M. K. Kim, I. K. Chung, Loss of RNA-binding protein HuR facilitates cellular senescence through posttranscriptional regulation of TIN2 mRNA. Nucleic Acids Res 46, 4271–4285 (2018).

61. W. Gong, S. Zhang, YB1 participated in regulating mitochondrial activity through RNA replacement. Front Oncol 13, 1145379 (2023).

62. T. Hofer, C. Badouard, E. Bajak, J. L. Ravanat, A. Mattsson, I. A. Cotgreave, Hydrogen peroxide causes greater oxidation in cellular RNA than in DNA. Biol Chem 386, 333–337 (2005).

63. Z. Ilic, A. R. Saxena, S. Periasamy, D. R. Crawford, Control (Native) and oxidized (DeMP) mitochondrial RNA are proinflammatory regulators in human. Free Radic Biol Med 143, 62–69 (2019).

64. H. Mirzaei, F. Regnier, Protein-RNA cross-linking in the ribosomes of yeast under oxidative stress. J Proteome Res 5, 3249–3259 (2006).

65. H. Wang, X. Wei, L. Liu, J. Zhang, H. Li, Suppression of A-to-I RNA-editing enzyme ADAR1 sensitizes hepatocellular carcinoma cells to oxidative stress through regulating Keap1/Nrf2 pathway. Exp Hematol Oncol 13, 30 (2024).

66. J. Yoon et al., Mitochondrial double-stranded RNAs as a pivotal mediator in the pathogenesis of Sjӧgren’s syndrome. Mol Ther Nucleic Acids 30, 257–269 (2022).

67. M. P. Murphy, L. A. J. O’Neill, A break in mitochondrial endosymbiosis as a basis for inflammatory diseases. Nature 626, 271–279 (2024).

68. N. S. Gokhale et al., Cellular RNA interacts with MAVS to promote antiviral signaling. Science 386, eadl0429 (2024).

69. B. Li, C. N. Dewey, RSEM: accurate transcript quantification from RNA-Seq data with or without a reference genome. BMC Bioinformatics 12, 323 (2011).

70. A. Dobin et al., STAR: ultrafast universal RNA-seq aligner. Bioinformatics 29, 15–21 (2013).

71. M. I. Love, W. Huber, S. Anders, Moderated estimation of fold change and dispersion for RNA-seq data with DESeq2. Genome Biol 15, 550 (2014).

72. G. Yu, L. G. Wang, Y. Han, Q. Y. He, clusterProfiler: an R package for comparing biological themes among gene clusters. OMICS 16, 284–287 (2012).

73. J. Joung et al., Genome-scale CRISPR-Cas9 knockout and transcriptional activation screening. Nat Protoc 12, 828–863 (2017).

74. C. Galaxy, The Galaxy platform for accessible, reproducible, and collaborative data analyses: 2024 update. Nucleic Acids Res 52, W83–W94 (2024).

75. D. A. Clayton, G. S. Shadel, Isolation of mitochondria from tissue culture cells. Cold Spring Harb Protoc 2014, pdb prot080002 (2014).

76. S. M. Blue et al., Transcriptome-wide identification of RNA-binding protein binding sites using seCLIP-seq. Nat Protoc 17, 1223–1265 (2022).

77. E. L. Van Nostrand et al., Robust transcriptome-wide discovery of RNA-binding protein binding sites with enhanced CLIP (eCLIP). Nat Methods 13, 508–514 (2016).

78. S. Heinz et al., Simple combinations of lineage-determining transcription factors prime cis-regulatory elements required for macrophage and B cell identities. Mol Cell 38, 576–589 (2010).

79. D. Y. Wu, D. Bittencourt, M. R. Stallcup, K. D. Siegmund, Identifying differential transcription factor binding in ChIP-seq. Front Genet 6, 169 (2015).

